# A VersaTile approach to reprogram the specificity of the R2-type tailocin towards different serotypes of *Escherichia coli* and *Klebsiella pneumoniae*

**DOI:** 10.1101/2024.10.29.620980

**Authors:** Dorien Dams, Célia Pas, Agnieszka Latka, Zuzanna Drulis-Kawa, Lars Fieseler, Yves Briers

## Abstract

Phage tail-like bacteriocins, or tailocins, provide a competitive advantage to producer cells by killing closely related bacteria. Morphologically similar to headless phages, their narrow target specificity is determined by receptor-binding proteins (RBPs). While RBP engineering has been used to alter the host range of a selected R2 tailocin from *Pseudomonas aeruginosa*, the process is labor-intensive, limiting broader application. We introduce a VersaTile-driven R2 tailocin engineering platform to scale up RBP grafting. This platform achieved three key milestones: (1) engineering R2 tailocins specific to *Escherichia coli* serogroups O26, O103, O104, O111, O145, O146 and O157; (2) reprogramming R2 tailocins to target for the first time capsule and a new species, specifically the capsular serotype K1 of *E. coli* and K11 and K63 of *Klebsiella pneumoniae*; (3) creating the first bivalent tailocin with a branched RBP and cross-species activity, effective against both *E. coli* K1 and *K. pneumoniae* K11. Over 90% of engineered tailocins were effective, with clear pathways for further optimization identified.

**Importance:** While tailocin engineering is a proven and promising concept, the current engineering approach lacks scalability, limiting a vast exploration. This study advances tailocin engineering by increasing its throughput. Implementing a scaled up approach, we have shown the flexibility of the R2 tailocin scaffold to accommodate diverse receptor-binding domains, expanding its functionality to target a new type of receptor (capsule) and a previously untargeted species. In addition, functional tailocins with branched receptor-binding proteins portraying dual, cross-genus activity were produced. This work lays the groundwork for a scalable platform for the development of engineered tailocins, marking an important step towards making R2 tailocins a practical therapeutic tool for targeted bacterial infections.

## Introduction

Bacterial infections have re-emerged as a significant threat to public health in the 21st century. Given the global spread of pathogens with increasingly accumulated antibiotic resistance mechanisms, the scientific community is compelled to develop alternative treatment therapies (1, 2). A potential class of novel antibacterials is phage tail-like bacteriocins (PTLBs), also referred to as tailocins (3). Tailocins are high-molecular-weight (>10^6^ Da), non-replicative protein complexes that share structural similarities with tailed bacteriophages but lack a capsid and a genome. This distinguishes them from phages, which are replicative entities (4). Upon induction of the SOS response, tailocins are intracellularly synthesized by bacteria and subsequently released to kill competing bacteria within the same niche, providing resistant sister cells a selective advantage for colonization. Morphologically and functionally, tailocins can be classified into R- and F-type tailocins (4).

R-type tailocins resemble the contractile tail of a T-even phage with a myovirus morphology and consist of an inner tubular core surrounded by an external sheath, as well as a baseplate to which receptor-binding proteins (RBPs) are anchored (5). R-type tailocins kill their target cell by binding to the cell surface, inducing structural changes in the baseplate of the tailocin. Subsequently, these changes trigger external sheath contraction, driving the inner tubular core to penetrate the bacterial cell envelope. As R2-type tailocins lack a capsid, they form a direct channel between the interior of the cell and the external environment, leading to uncontrolled ion leakage and collapse of the membrane potential. As a consequence, membrane dissipation compromises essential cellular processes, ultimately leading to cell death (4, 6). F-type tailocins, in contrast, resemble the non-contractile tail of a lambda phage, having a siphovirus morphology with a different but unknown killing mechanism. Some bacteria, especially *Pseudomonas aeruginosa*, produce both R- and F-type tailocins under the control of the same regulatory genes and are released via the same lysis genes (4, 7).

Similar to phages, the host spectrum of tailocins is determined by its receptor-binding protein (RBP), which recognizes and binds a receptor located on the bacterial surface (*e.g.*, polysaccharides or outer membrane proteins). RBPs are typically homotrimers with a modular structure comprising a conserved N-terminal domain responsible for anchoring to the phage tail or tailocin baseplate, and a variable C-terminal receptor-binding domain (RBD) responsible for host surface receptor recognition (8). The high amino acid sequence variability observed in the RBD of RBPs mostly enables precise subspecies-specific recognition, leading to a distinct killing spectrum for each tailocin. The subspecies target specificity of tailocins also implies that the target pathogen should be well known prior to successful tailocin application, or that their cocktails must cover a variation of targeted strains, similar to the *sur mesure* and *prêt-à-porter* paradigms in the phage therapy field (9). The use of narrow-spectrum antibacterials is increasingly considered beneficial due to their microbiota-friendly nature and the eminent role of a balanced microbiome for human health (10). There is thus a need to create a large collection of custom tailocins for therapeutic purposes. However, tailocins are presently only identified in a handful of species (11).

Recent advancements in RBP engineering have highlighted the possibility of adjusting the host range of phages and tailocins by substituting complete RBPs or RBDs with their equivalents originating from different phages, prophages, or tailocins, into conserved viral or tailocin scaffolds (12–16). The R2 tailocin produced by *Pseudomonas aeruginosa*, also named R2 pyocin, is the best-studied tailocin scaffold and has demonstrated its capacity to accept heterologous C-terminal RBDs with both tail fiber and tailspike characteristics originating from different phages (4). As such, the antimicrobial range of the R2 tailocin has been successfully reprogrammed across species borders towards *Yersinia pestis* (17), *Escherichia coli* (17–20), *Salmonella enterica* (21), and *Campylobacter jejuni* (22). To become a broadly applicable solution of customized, narrow-spectrum antibacterials either for *sur mesure* or *prêt-à-porter* applications, the R2 tailocin engineering approach should be turned into a convenient platform for the rapid creation of tailor-made tailocins. Until now, studies are limited to only one to a few RBPs, underscoring that tailocin RBP engineering has not yet matured into an easily scalable approach. To produce modified R2 tailocins, the *P. aeruginosa* host strain PAO1 Δ*prf15*, encoding the full operon for the biosynthesis of the R2 tailocin but deficient in its RBP gene is complemented *in trans* with an expression vector producing a chimeric RBP. This chimeric RBP is composed of the native R2 N-terminal anchor to attach the chimeric RBP to the tailocin particle and a C-terminal RBD recognizing the receptor of interest. In previous work, tailocin RBP engineering relies on restriction-ligation reactions with traditional type IIp restriction enzymes (REs) to create RBP fusions in expression vectors, which is a cumbersome and tedious approach.

This study aims to introduce a more flexible and convenient approach as an initial step to scale up RBP grafting in generalized scaffolds by implementing the VersaTile technique (23). VersaTile is a combinatorial DNA assembly technique that is convenient for the assembly of non-homologous sequences of modular proteins such as RBPs. It implies two steps: (1) Creating a repository of so-called ‘tiles’, consisting of a DNA sequence-of-interest flanked with location-specific position tags; (2) Assembling all tiles in the appropriate vector in a one-step restriction-ligation reaction using only one type of IIs RE (23). Moreover, all tiles can be easily reused to assemble different constructs in various vectors and thus many scaffolds, allowing a better scalable plug-and-play Lego-like approach.

As a proof of concept of a tailocin platform with pluggable RBPs, we have grafted RBPs recognizing the O-antigen of *E. coli*, and the K-antigen (or capsule) of *E. coli* and *Klebsiella pneumoniae* in the R2 tailocin scaffold. Both the capsule and the *Klebsiella* species were earlier not yet targeted by engineered tailocins. The variety of O- and K-antigens in *E. coli* and *K. pneumoniae* generally correlates well with the diversity of RBPs produced by phages infecting *E. coli* and *K. pneumoniae*, respectively. Numerous RBPs contain a specific polysaccharide-depolymerizing domain cleaving LPS O-antigen or degrading capsule. Whereas the natural R2 tailocin of *P. aeruginosa* contains a single RBP, we demonstrate that a functional bivalent tailocin with a more complex, branched RBP structure targeting two receptors from different species can be created.

## Results

### A plug-and-play R2 tailocin engineering platform

The VersaTile DNA assembly technique was implemented into the RBP engineering of the tailocin engineering platform, allowing rapid and straightforward engineering of chimeric RBPs. The engineered RBP constructs consist of two types of tiles. First, the anchor tile originates from the native R2 tailocin and is flanked by two codon-long position tags P_start_ and P_mid_. Second, the receptor-binding domain (RBD) tile originates from a wide range of Escherichia and Klebsiella phage RBPs targeting O-antigen or K-antigen (capsule) and is flanked by position tags P_mid_ and P_end_. These position tags enable easy and standardized recombination of the two types of tiles in the correct order into the VersaTile-adjusted *P. aeruginosa*-*E. coli* shuttle expression vector pVTD29 used for *in trans* expression of the chimeric RBP (**FIG 1**).

**FIG 1.**
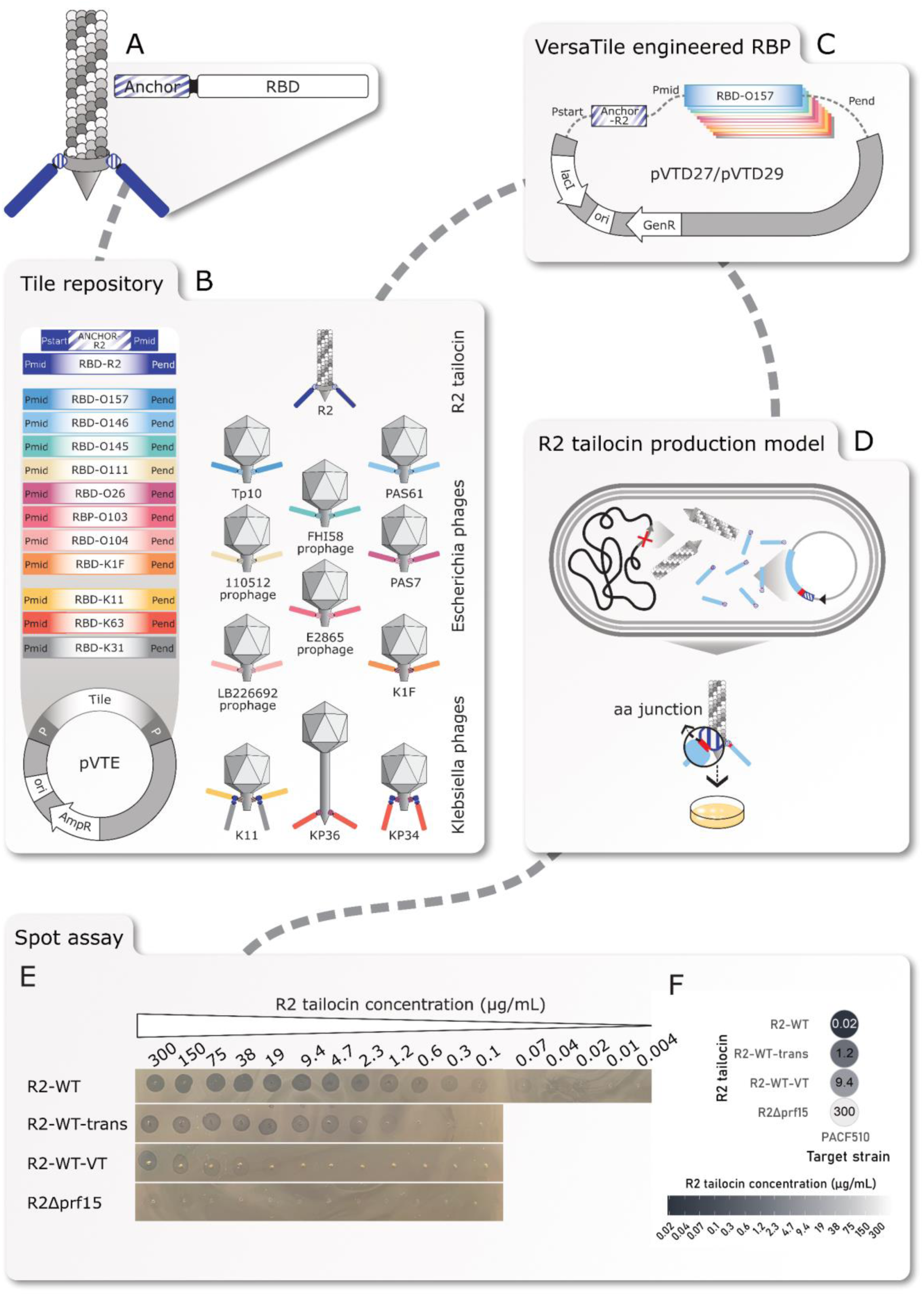
Pipeline for tailocin engineering and production. (**A**) Modular build-up of the R2 tailocin receptor-binding protein (RBP). (**B**) Establishment of a tile repository consisting of anchor and receptor-binding domain (RBD) tiles sourced from the R2 tailocin, and phages infecting *Escherichia coli* or *Klebsiella pneumoniae*, cloned in the pVTE entry vector using VersaTile cloning. (**C**) The use of VersaTile assembly to combine the anchor and RBD tiles in a predefined order in the expression vector. (**D**) The production of engineered tailocins in the producer host PAO1 Δ*prf15* upon induction of the SOS response with mitomycin C to produce the RBP-deficient mutant R2Δ*prf15*, and *in trans* expression of the chimeric RBP with IPTG. (**E**) Results of the spot assay with visible zones at different concentrations of the spotted R2 wild-type (R2-WT) and its derivates, with the *in trans* expression of the chimeric RBP (R2-WT-trans), the *in trans* expression of the VersaTile assembled RBP (R2-WT-VT), and the expression of the deficient RBP (R2Δ*prf15*) alone. (**F**) Overview of the results of the spot assay for all R2 wild-type tailocin derivatives. Lower concentrations indicate a higher R2 tailocin activity, and the lowest concentrations at which visible clearance was observed with the naked eye on a bacterial lawn of the *P. aeruginosa* target strain CF510 are displayed.

Three positive controls were implemented: (1) The native R2 tailocin (R2-WT) in which the RBP was expressed from the *P. aeruginosa* PAO1 genome. (2) The native R2 tailocin with *in trans* expression of the RBP (R2-WT-trans). Here, the tailocin scaffold is produced by *P. aeruginosa* strain PAO1 Δ*prf15*, a host with a deficiency in gene *prf15* encoding the RBP. The wild-type RBP is provided *in trans* using expression vector pVTD27, without a *lac* repressor, or pVTD29, with a *lac* repressor. (3) The native R2 tailocin with *in trans* expression of the VersaTile assembled RBP (R2-WT-VT). The RBP coding sequence has been reconstructed with the VersaTile DNA assembly technique which introduces (similar to traditional Type IIp restriction enzymes) a six-nucleotide junction between the anchor and RBD. Additionally, a negative control was co-expressed, namely the R2 tailocin lacking the RBP, produced by *P. aeruginosa* Δ*prf15* (R2Δ*prf15*).

### Validation of the R2 tailocin production platform through the production of four R2 tailocin controls

All R2 tailocin wild-type derivatives mentioned above were purified using the ammonium sulphate (AS) precipitation method. A two-fold dilution series starting from 300 µg/mL was tested. First, the native R2 tailocin (R2-WT) exhibited clearance at a concentration as low as 0.02 µg/mL (**FIG 1**, e and f). Transparency of the spots decreases with lower R2 tailocin concentrations. Next, R2-WT-trans and R2-WT-VT were evaluated to assess the *in trans* expression efficiency of the RBP using the VersaTile adapted vector, and the effect of the six-nucleotide junction between the anchor tile and the RBD tile. Here, opaque spots were observed up to a concentration of 1.2 µg/mL for R2-WT-trans and 9.4 µg/mL for R2-WT-VT. This suggests a 60-fold lower efficiency when the R2 tailocin is expressed *in trans* along with an additional 8-fold reduction in tailocin activity due to the presence of the junction. Important to note is that R2Δ*prf15* also resulted in an opaque spot in the spotting assay at a high concentration of 300 µg/µL, which may be attributed to F-type tailocins produced under the same regulatory genes of the SOS response triggered by mitomycin C (7).

### Confirmation of R2-WT, R2-WT-trans, and R2-WT-VT activities by quantitative survival and growth inhibition assays

R2-WT, R2-WT-trans and R2-WT-VT showed a dose-dependent killing (**FIG 2**, a), starting from 0.8, 1.6, and 50 µg/mL, respectively. A dose of 100 µg/mL of R2-WT, R2-WT-trans and R2-WT-VT reduced the cell count with 6.6 ± 2.0, 4.5 ± 0.6 and 2.5 ± 1.4 log, respectively (**TABLE 1**).

**FIG 2.**
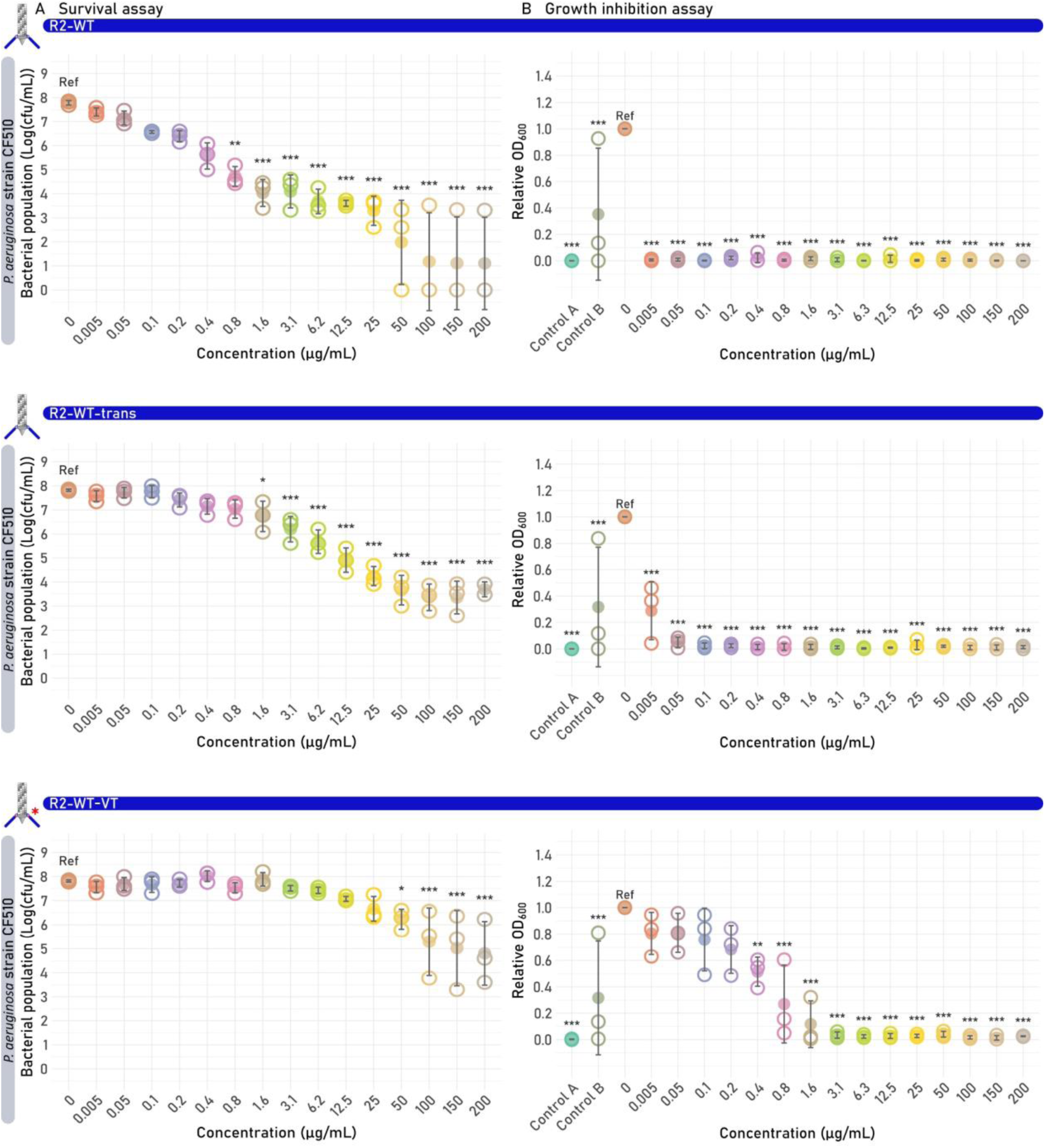
Results of the survival and growth inhibition assays of the R2 wild-type tailocin derivatives R2-WT, R2-trans, and R2-VT. Each R2 tailocin was tested on the susceptible target strain *P. aeruginosa* CF510. Significant differences are shown by asterisks (*p < 0.05; **p < 0.005, ***p < 0.001). (**A**) Survival assay. One plot is shown for each R2 tailocin construct, showing the bacterial colony count in function of the concentration of the added R2 tailocin (derivative). The value of each biological replicate is displayed using open circles and the mean values are shown as full circles. (**B**) Growth inhibition assay results at 8 h are shown per R2 tailocin construct. For each R2 tailocin concentration. The relative turbidity of each biological replicate is displayed using open circles, and the mean relative turbidity is shown as full circles. Two additional controls were performed, one containing R2 tailocin but lacking the bacterial strain (Control A) and one containing the R2 tailocins particle lacking the RBP (R2Δ*prf15*) (Control B). Both controls were added at the highest available R2 tailocin concentration, which was 200 µg/mL for all R2 wild-type tailocin derivatives.

**TABLE 1.**
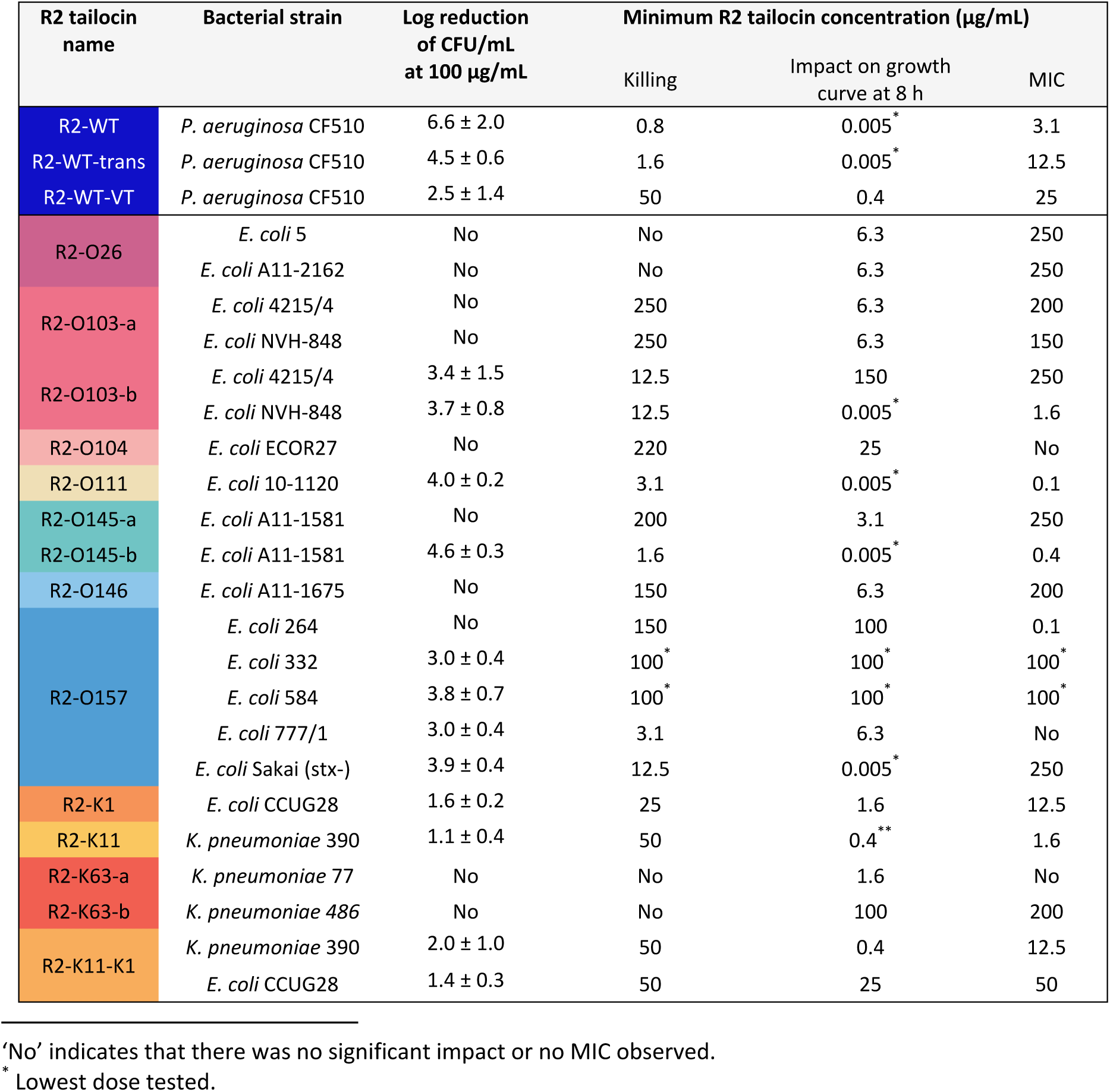
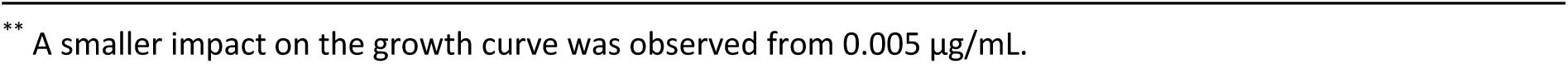
Overview of all survival and growth inhibition assays of the wild-type and engineered R2 tailocins. Four values are listed per R2 tailocin (derivative) obtained for their corresponding strain(s) tested: (1) Log reduction of the cell count (log(CFU/mL)) at an R2 tailocin concentration of 100 µg/mL in the survival assay; (2) The lowest R2 tailocin concentration giving a significant (p < 0.05) difference in bacterial count compared to the untreated sample in the survival assay; (3) The lowest R2 tailocin concentration giving a significant (p < 0.05) impact on the growth curve at time point 8 h compared to the untreated sample in the growth inhibition assay; (4) The minimum inhibitory concentration (MIC) at which > 90% inhibition of the growth curve was observed after 24 h.

| R2 tailocin name | Bacterial strain | Log reduction of CFU/mL at 100 μg/mL | Minimum R2 tailocin concentration (μg/mL) | | |
| --- | --- | --- | --- | --- | --- |
|  |  |  | Killing | Impact on growth curve at 8 h | MIC |
| R2-WT | *P. aeruginosa* CF510 | 6.6 ± 2.0 | 0.8 | 0.005* | 3.1 |
| R2-WT-trans | *P. aeruginosa* CF510 | 4.5 ± 0.6 | 1.6 | 0.005* | 12.5 |
| R2-WT-VT | *P. aeruginosa* CF510 | 2.5 ± 1.4 | 50 | 0.4 | 25 |
| R2-O26 | *E. coli* 5 | No | No | 6.3 | 250 |
|  | *E. coli* A11-2162 | No | No | 6.3 | 250 |
| R2-O103-a | *E. coli* 4215/4 | No | 250 | 6.3 | 200 |
|  | *E. coli* NVH-848 | No | 250 | 6.3 | 150 |
| R2-O103-b | *E. coli* 4215/4 | 3.4 ± 1.5 | 12.5 | 150 | 250 |
| | *E. coli* NVH-848 | 3.7 ± 0.8 | 12.5 | 0.005* | 1.6 |
| R2-O104 | *E. coli* ECOR27 | No | 220 | 25 | No |
| R2-O111 | *E. coli* 10-1120 | 4.0 ± 0.2 | 3.1 | 0.005* | 0.1 |
| R2-O145-a | *E. coli* A11-1581 | No | 200 | 3.1 | 250 |
| R2-O145-b | *E. coli* A11-1581 | 4.6 ± 0.3 | 1.6 | 0.005* | 0.4 |
| R2-O146 | *E. coli* A11-1675 | No | 150 | 6.3 | 200 |
| R2-O157 | *E. coli* 264 | No | 150 | 100 | 0.1 |
| | *E. coli* 332 | 3.0 ± 0.4 | 100* | 100* | 100* |
| | *E. coli* 584 | 3.8 ± 0.7 | 100* | 100* | 100* |
| | *E. coli* 777/1 | 3.0 ± 0.4 | 3.1 | 6.3 | No |
| | *E. coli* Sakai (stx-) | 3.9 ± 0.4 | 12.5 | 0.005* | 250 |
| R2-K1 | *E. coli* CCUG28 | 1.6 ± 0.2 | 25 | 1.6 | 12.5 |
| R2-K11 | *K. pneumoniae* 390 | 1.1 ± 0.4 | 50 | 0.4** | 1.6 |
| R2-K63-a | *K. pneumoniae* 77 | No | No | 1.6 | No |
| R2-K63-b | *K. pneumoniae* 486 | No | No | 100 | 200 |
| R2-K11-K1 | *K. pneumoniae* 390 | 2.0 ± 1.0 | 50 | 0.4 | 12.5 |
| | *E. coli* CCUG28 | 1.4 ± 0.3 | 50 | 25 | 50 |

'No' indicates that there was no significant impact or no MIC observed.
\* Lowest dose tested.
\*\* A smaller impact on the growth curve was observed from 0.005 µg/mL.

In the growth inhibition assay starting from a low inoculum (± 10^5^ CFU/mL), growth was monitored over a period of 24 h upon addition of the tailocin. Growth was evaluated at two different time points: (I) After 8 h, the difference in optical density between various concentrations of the engineered R2 tailocin and the untreated sample was evaluated. (II) After 24 h, the minimum inhibitory concentration (MIC) for full inhibition was investigated. R2-WT and R2-WT-trans impacted the growth curve at the lowest concentration tested (0.005 µg/ml) at 8 h, while R2-WT-VT showed inhibition starting from 0.4 µg/ml (**FIG 2**, b). The MIC after 24 h was 3.1, 12.5, and 50 µg/mL, for R2-WT, R2-WT-trans, and R2-WT-VT, respectively (**TABLE 1**; **Supplementary FIG S1**). However, it should be noted that while R2Δ*prf15* (control B; high dose of 200 µg/mL) showed no activity in the survival assay, a significant impact was observed on the growth curve, possibly attributable to the simultaneous production of F-type tailocins.

Altogether, these data demonstrate that functional tailocins are successfully produced using the VersaTile-compatible tailocin engineering platform, emphasizing its potential for rapid, convenient, and high-throughput screening. Yet, reductions in efficacy must be taken into account for *in trans* expression, likely due to an imbalanced production (in amount or time) of the RBP versus the tailocin scaffold, impacting tailocin assembly and the proportion of correctly assembled tailocins. The two amino acid junction further impacts efficacy, at least for the evolutionary optimized native R2 tailocin of *P. aeruginosa*. Plausible causes are an inferior orientation of the RBP, a negatively impacted signal cascade linking cell binding and the eventual killing mechanism or differences in RBP expression yield.

### Construction of nine chimeric RBPs targeting *E. coli* serogroups O26, O103, O104, O111, O145, O146, and O157

The C-terminal RBDs targeting *E. coli* O-antigen originate from phages with a podovirus morphology, including the RBPs from Escherichia phage PAS7 (O103) belonging to the recently proposed genus *Cepavirus* (24), Escherichia phages PAS61 (O146) and O157 typing phage 10, or Tp10 for short (O157), belonging to the genus *Uetakevirus*. Other RBPs originate from prophages belonging to the *Lederberg* (O111, O145) and *Uetakevirus* (O26) genera that were identified within the genomes of bacterial strains with the desired serogroups, namely *E. coli* strains RM10386 (O26), 110512 (O111) and FHI58 (O145). Fusions between the N-terminal R2 anchor and the C-terminal RBDs from the mentioned *E. coli* (pro)phages using VersaTile were made to retarget the bactericidal spectrum of the native R2 tailocin towards specific *E. coli* O-antigen serogroups. These engineered R2 tailocins were produced and purified using high-speed centrifugation with protein yields ranging from 439 to 857 µg/mL (**Supplementary FIG S2**). For two engineered R2 tailocins containing RBDs against serogroups O103 and O145, an alternative shuttle expression vector (pVTD27) was used for comparison. pVTD27 is identical to pVTD29 but does not contain a *lac* repressor (*lacI* gene). The *lac* repressor prevents leaky expression from the used *lac* promotor, which is induced by isopropyl ß-D-1-thiogalactopyranoside (IPTG), generally lowering the expression level of the RBP gene prior to induction.

### All engineered R2 tailocins targeting *E. coli* O-antigens are bactericidal against at least one susceptible strain with highly variable efficacy

In a spot assay zones of clearance were observed on at least one susceptible strain for all engineered R2 tailocins targeting a specific *E. coli* O-antigen (**FIG 3**). The best-performing engineered R2 tailocins are R2-O145-b, R2-O103-b, and R2-O157, and R2-O111, showing clearance at 0.07 µg/mL, 0.6 µg/mL, 0.6 µg/mL and 2.3 µg/mL (corresponding to 3.5, 30, 30 and 115 × higher concentrations than the average R2-WT). R2-O26, R2-O104, and R2-O146 were active but remarkably higher doses (75-300 µg/mL) were needed to be visible as a spot. Some strains were insensitive for the corresponding engineered tailocins. These observations and the variability therein indicate that the R2 tailocin scaffold is remarkably amenable to accept a diversity of new RBDs. It should be emphasized that the newly grafted RBDs in this study have a tailspike morphology in contrast to the native tail fiber structure. Nevertheless, the inferior activity compared to the native evolutionarily optimized R2 tailocin of *P. aeruginosa* shows that there is further technical optimization potential (delineation, shuttle vector, different RBD source) as seen for the two variants of R2-O103 and R2-O145. In addition, a horizontal transfer event in nature is accompanied with adaptive evolution, particularly surrounding the newly formed junction site to optimize the functioning of the new chimer. It is plausible that mutagenesis around the junction site will further enhance the antibacterial effect.

**FIG 3.**
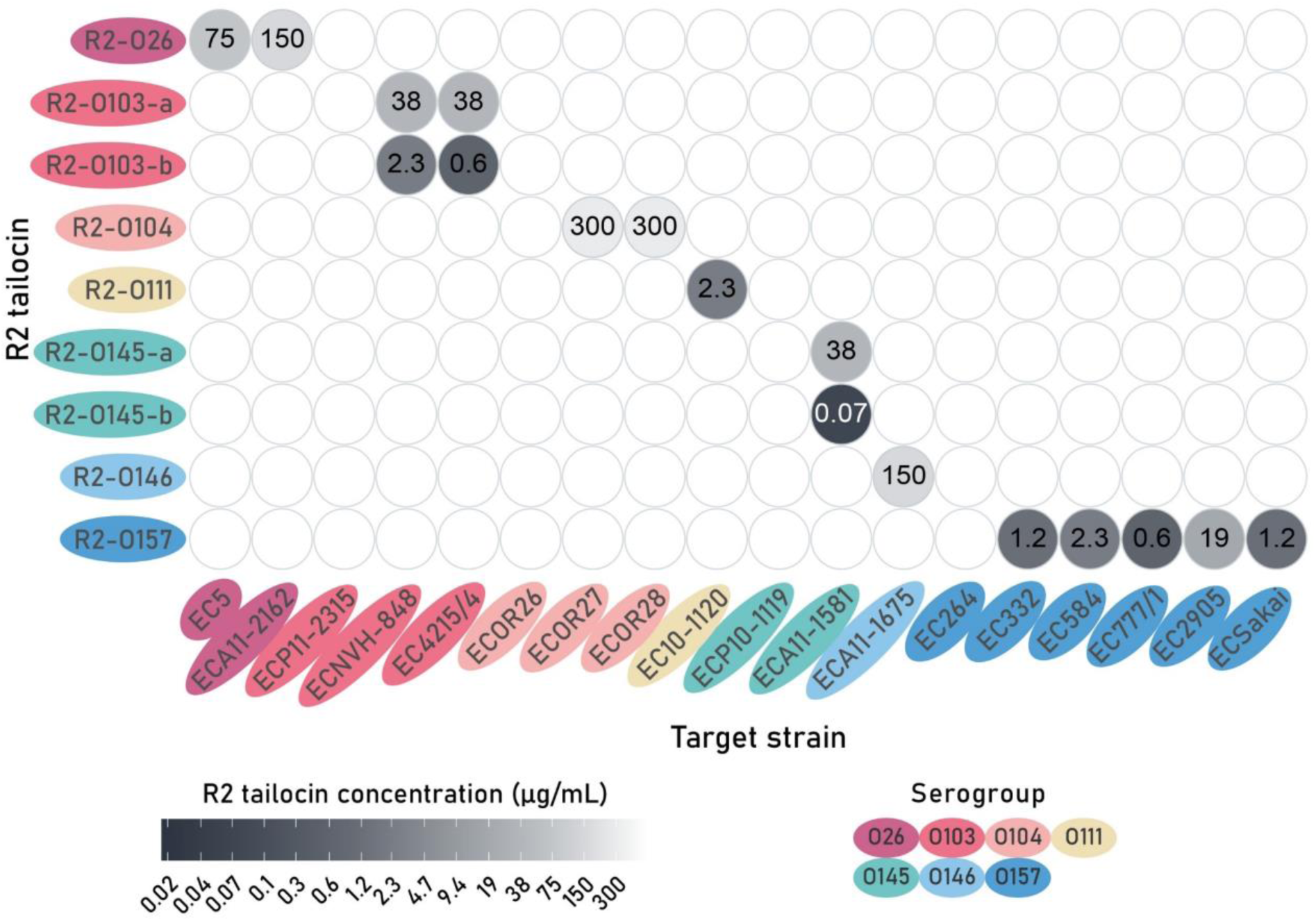
Spot assay of all engineered R2 tailocins tested against all available *Escherichia coli* strains of different serogroups. The lowest concentrations at which clearance was observed on bacterial lawns of *E. coli* target strains are displayed. Lower concentrations indicate a higher R2 tailocin activity. The engineered R2 tailocins and target strains are organized and colored according to O-antigen serogroups of the phage host donating the RBD and the target *E. coli* strain.

The strains susceptible to a specific tailocin in the spot assay also gave variable outcomes in the survival and growth inhibition assays. R2-O145-b, R2-O111 and R2-O157 caused the highest reduction in bacterial cell count (between 3 and 4.6 log across the susceptible *E. coli* strains tested) in the survival assay, with 1.6, 3.1 and 3.1 µg/mL as the lowest concentration with significant reduction, followed by R2-O103-b (12.5 µg/mL; 3.4-3.7 log) (**TABLE 1**; **FIG 4**). Other engineered tailocins required doses higher than 150 µg/mL and for R2-O26, no significant reduction in cell count could be observed. The growth inhibition assay evaluated at 8 h was the most sensitive metric to identify (partially) functional tailocins, with a significant impact on the growth curve at 8 h for all engineered tailocins (**TABLE 1**; **FIG 4**; **Supplementary FIG S1)**. Generally, higher doses were needed to maintain a significant impact on the growth curve after 24 h, which can be explained by an incomplete eradication of the targeted bacteria or the growth of emerging resistant clones.

**FIG 4.**
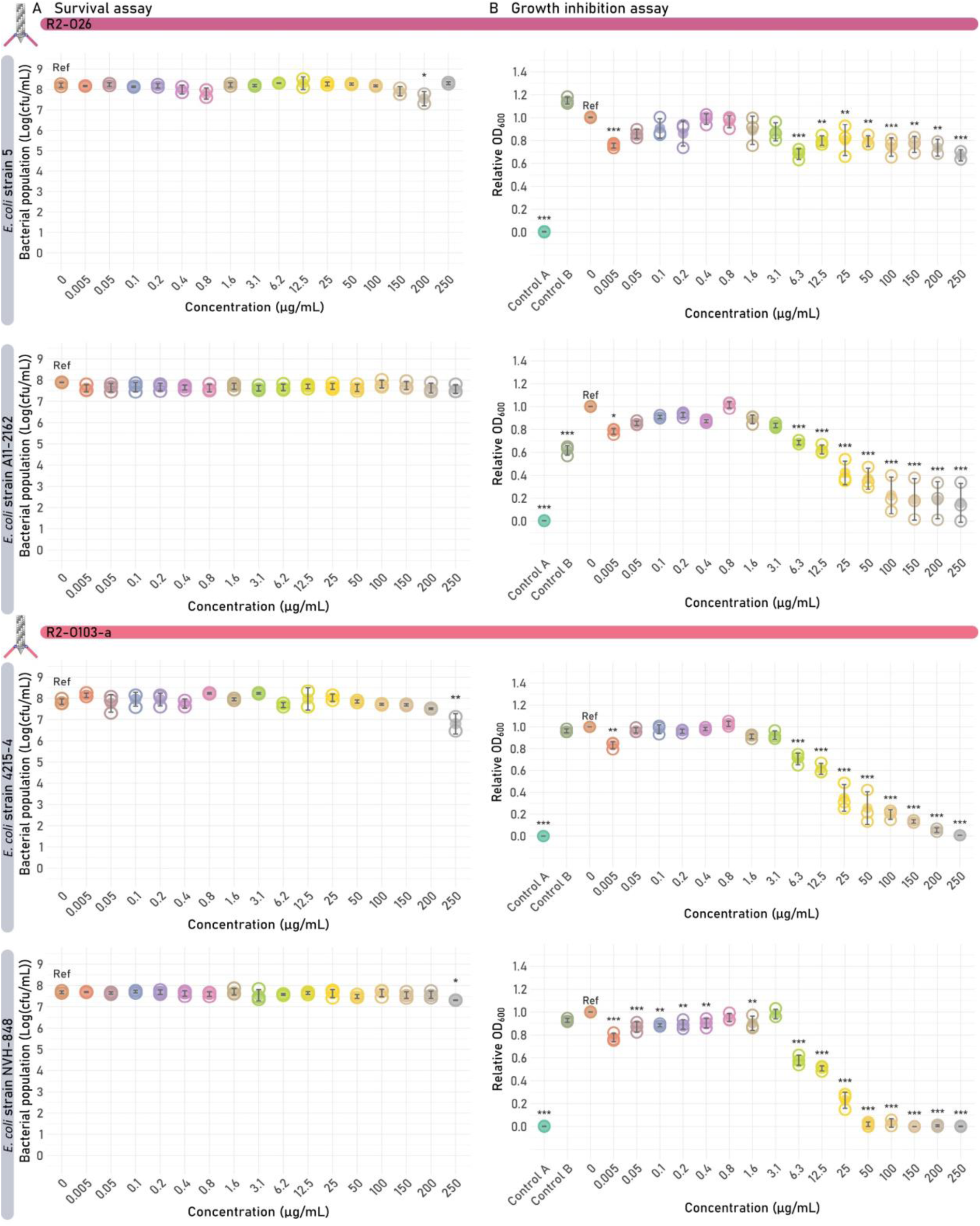
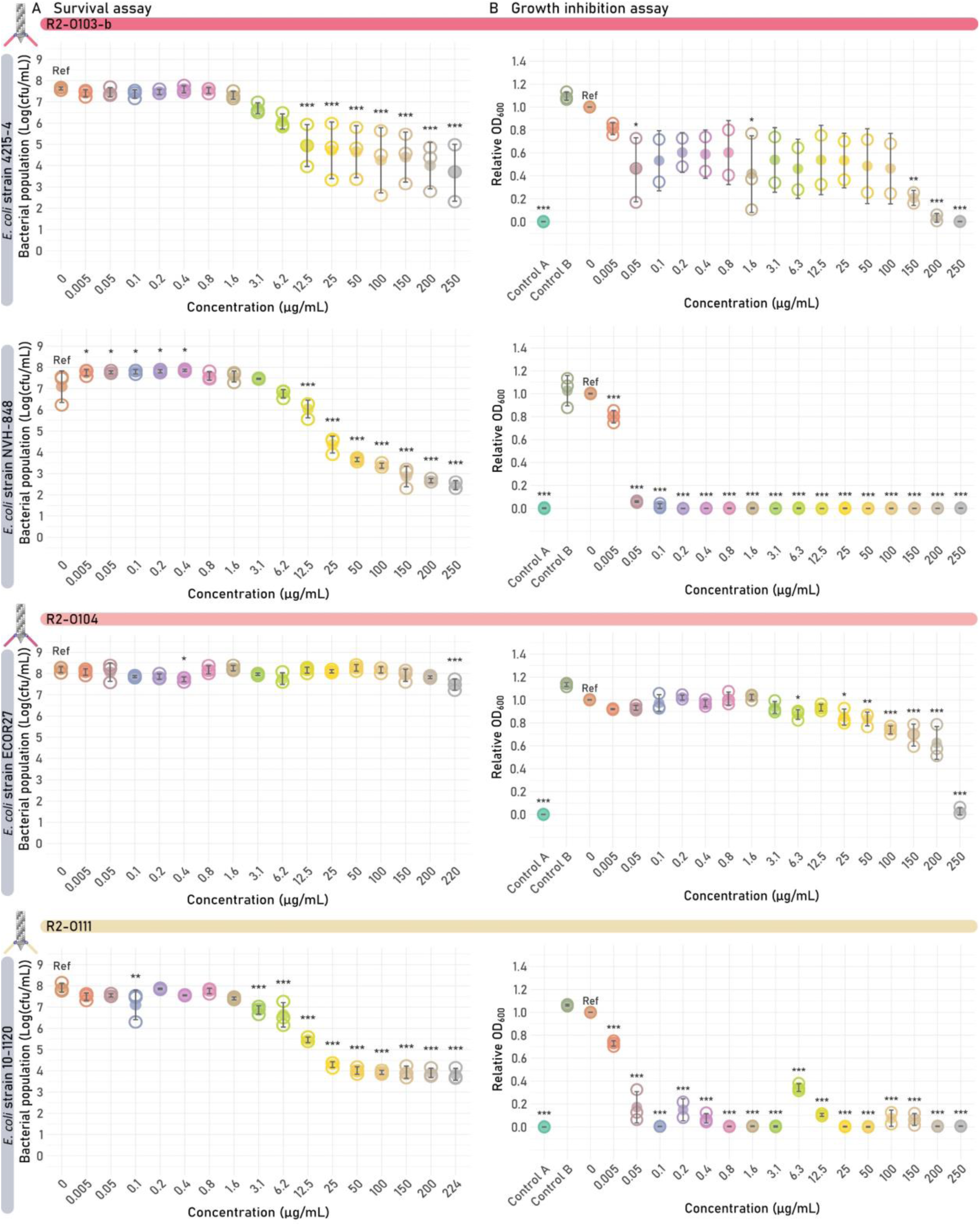
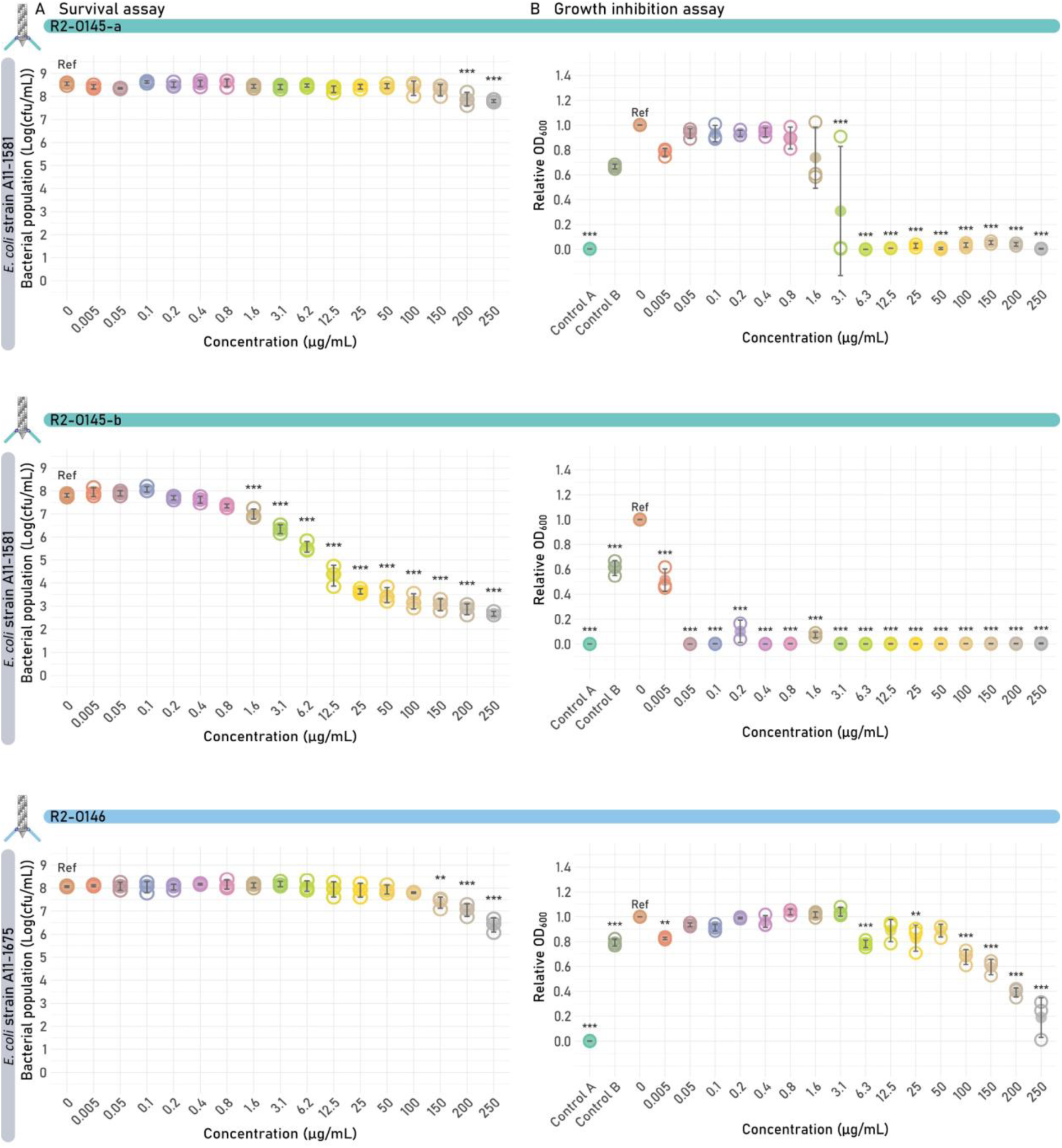
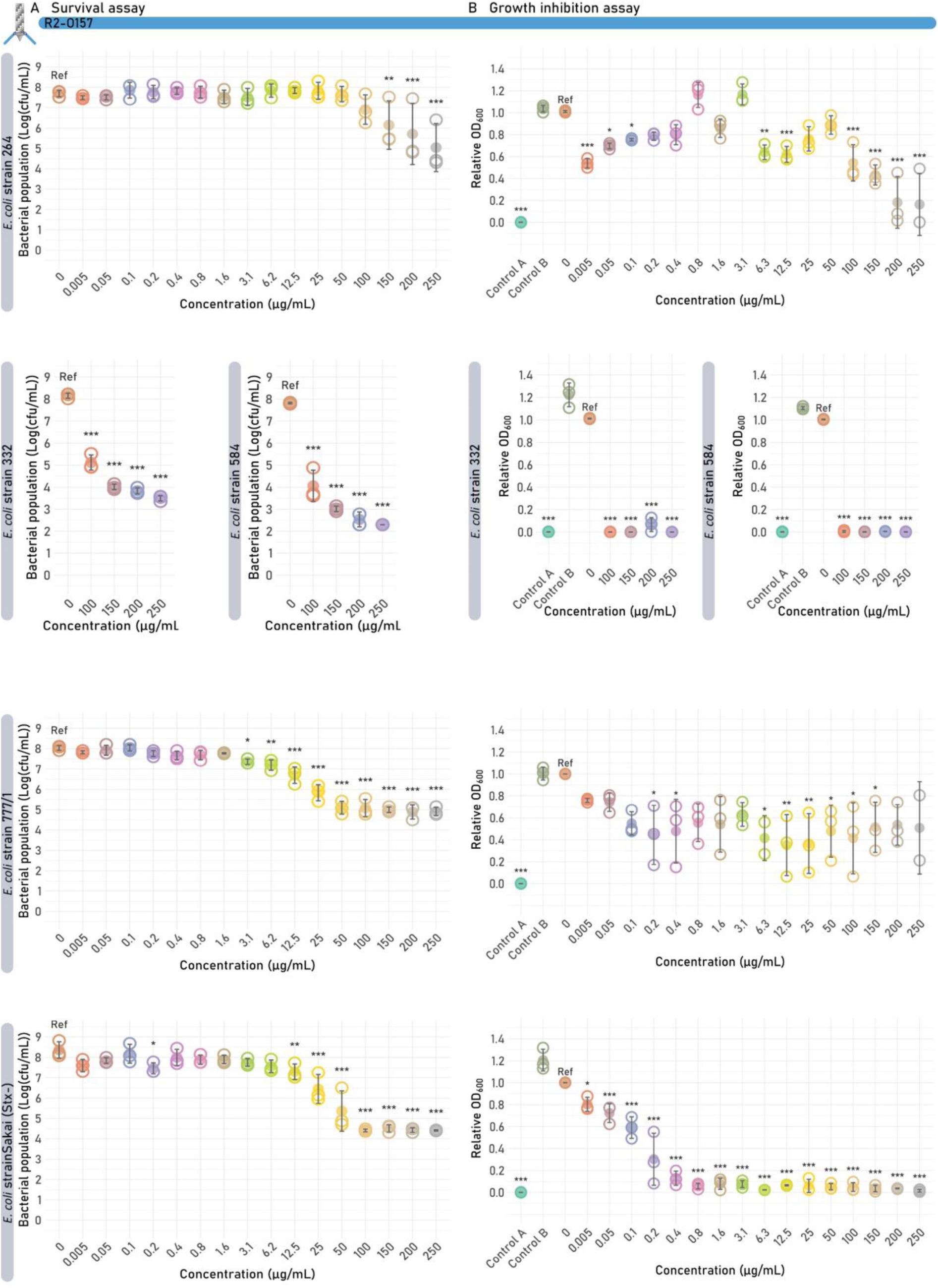
Survival and growth inhibition assays of the different engineered *Escherichia coli* O-antigen targeting R2 tailocins. Each R2 tailocin was tested on their susceptible *E. coli* target strains. Significant differences are shown by asterisks (*p < 0.05; **p < 0.005, ***p < 0.001). (**A**) Survival assay. One plot is shown for each R2 tailocin construct, showing the bacterial colony count in function of the concentration of the added R2 tailocin. The value of each biological replicate is displayed using open circles and the mean values are shown as filled circles. (**B**) Growth inhibition assay results at 8 h are shown per R2 tailocin construct. The relative OD_600_ of each biological replicate is displayed using open circles, and the mean relative OD_600_ is shown as full circles. Two additional controls were performed, one containing the R2 tailocin but without the bacterial strain (Control A) and one containing a receptor-binding protein (RBP) lacking mutant R2 tailocin particle (R2Δprf15) (Control B). Both controls were added at the highest available R2 tailocin concentration (220-250 µg/mL).

### Construction of five chimeric RBPs targeting *E. coli* capsular serotype K1 and *K. pneumoniae* capsular serotypes K11, K31, and K63

The tailocin engineering approach based on the VersaTile assembly technique allowed rapid screening of a large diversity of RBDs for the construction and identification of functionally engineered tailocins customized to an array of O-antigens. In addition to expanding the availability of *E. coli* serogroup-specific tailocins, this standardized approach was further applied to address another cell surface receptor and species. Many *E. coli* strains also possess a polysaccharide capsule (K-antigen) to increase their pathogenicity, with K1 as the most prevalent capsule serotype (25). Therefore, we selected the RBD of the K1 capsule-specific *E. coli* phage K1F (K1Fgp17) (podovirus morphology, genus *Kayfunavirus*). In addition, we sourced RBDs from several Klebsiella phages targeting capsular serotypes to engineer *Klebsiella-*specific R2 tailocins. Specifically, we selected RBDs from phages with podovirus (phages K11 and KP34) and siphovirus (phage KP36) morphologies. Both phages KP34 (*Drulisvirus*) and KP36 (*Webervirus*) possess a single RBD with 42.4% aa similarity (94% coverage) between them, recognizing the same capsular serotype K63, designated here as RBD-K63-a and RBD-K63-b, respectively. Phage K11 belongs to the genus *Przondovirus* and harbors a dual, branched RBP system, with each RBP having a different receptor specificity (capsular serotype K11 and K31) (26). All selected RBDs were fused to the N-terminal R2 anchor, and engineered tailocins were produced and purified by high-speed centrifugation, resulting in protein concentrations ranging from 273 to 470 µg/mL (**Supplementary FIG S2**).

### Engineered tailocins can target capsule as receptor against both *E. coli* and *K. pneumoniae*

R2-K1 was spotted against the panel of *E. coli* and *K. pneumoniae* strains (**FIG 5**). Besides capsule K1 producing *E. coli* strains CUGG28 and CAB1, R2-K1 also showed clearance on *E. coli* strain ECOR28 of serogroup O104, of which the capsular serotype was unknown. A blastn analysis was performed to search for similarities between the ECOR28 strain genome (GCF_002190675.1) with serotype-specific capsule biosynthesis proteins (25). This analysis revealed similarities (> 97% coverage) to the proteins encoded by genes *neuD, neuB, neuA* and *neuC* (respective UniProtKB accessions: Q46674, Q46675, A0A0D6H548, and Q47400) responsible for K1 capsule biosynthesis, with 59, 69, 48 and 59% aa identity to their respective protein sequences. However, no similarity was found to the remaining K1 capsule biosynthesis proteins encoded by genes *neuE* and *neuS* (UniProtKB accessions: Q47401 and Q47404). The *Klebsiella* strains were not susceptible for R2-K1.

**FIG 5.**
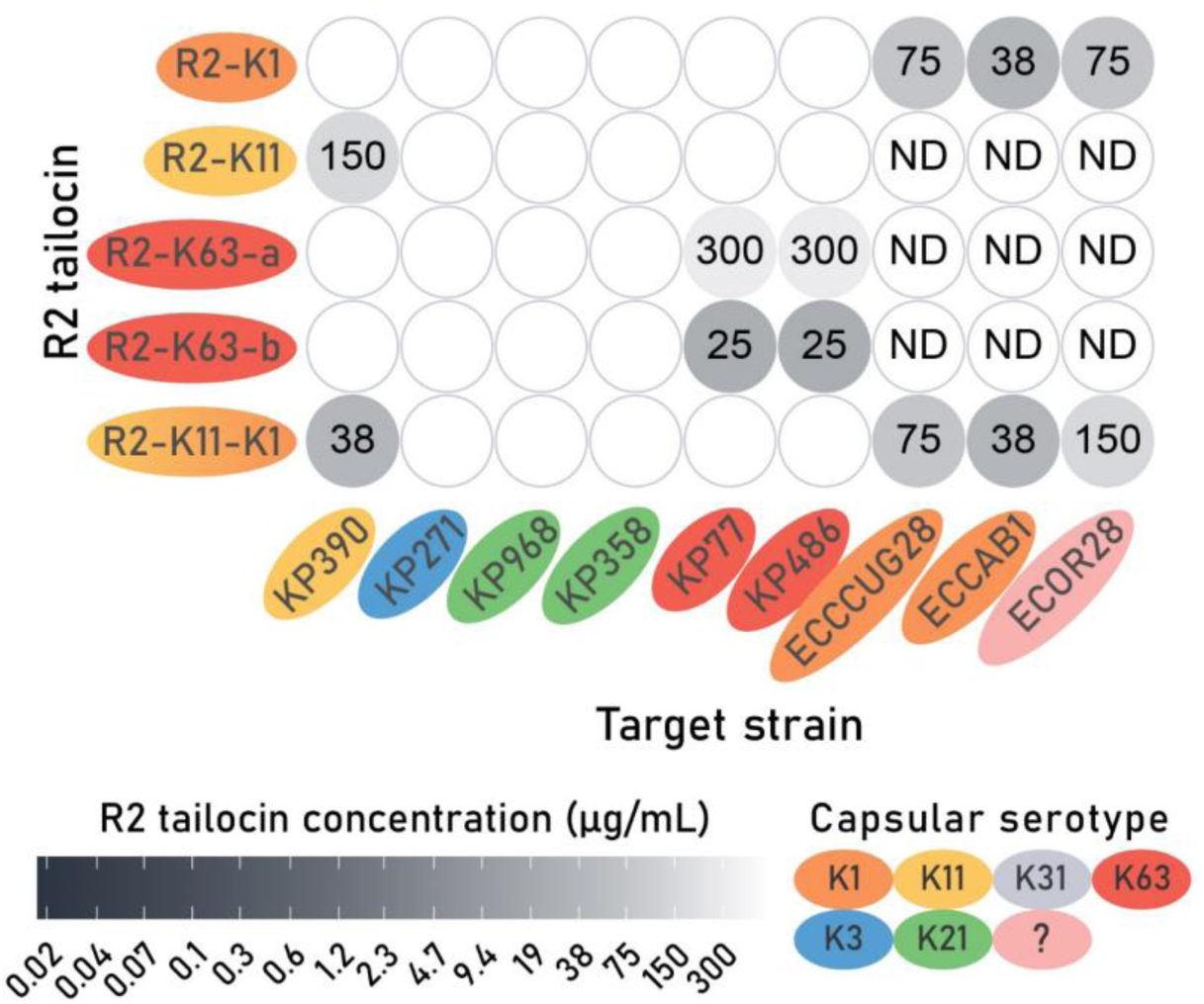
Spot assay of all engineered capsule-targeting R2 tailocins tested against *Klebsiella pneumoniae* and *Escherichia coli* capsular serotypes. The lowest concentrations at which clearance was observed on bacterial lawns of *K. pneumoniae* and *E. coli* target strains are displayed. Lower concentrations indicate a higher R2 tailocin activity. Engineered R2 tailocins that were not spotted against certain strains were indicated as not determined (ND). *E. coli* strain ECOR28 has an unknown capsular serotype as indicated by a question mark. The engineered R2 tailocins and target strains are organized and colored according to K-antigen serogroups of the phage host donating the RBD and the target *K. pneumoniae* or *E. coli* strain.

All Klebsiella-targeting tailocins were spotted against a panel of *K. pneumoniae* strains with different capsular serotypes. The observed susceptibilities fully agree with the host specificity of the corresponding phage sources (**FIG 5**). While in general the required minimum doses were higher compared to O-antigen targeting tailocins, R2-K63-b with the RBD of Klebsiella siphophage KP36 demonstrated the most effective clearance starting from a concentration of 25 µg/mL. In contrast, R2-K63-a derived from podophage KP34 requires a 12-fold higher dose. These observed differences demonstrate the value of an approach to rapidly construct and screen a diversity of RBDs in a standardized manner. Additionally, the RBD of K11gp43, targeting capsular serotype K31 was grafted into the R2 tailocin scaffold (R2-K31). Unfortunately, no spots were observed for R2-K31. However, only one *K. pneumoniae* strain with capsular serotype K31 was tested which could not be infected by phage K11. Therefore, this line of investigation was discontinued.

Survival and growth inhibition assays were conducted for tailocins that exhibited activity in the spot assay (**FIG 6**; **TABLE 1**). R2-K1 and R2-K11 showed a significant bactericidal effect against their susceptible host, starting from a concentration of 25 and 50 µg/mL, respectively. At a R2 tailocin concentration of 100 µg/mL, a bactericidal effect of 1.6 and 1.1 log(CFU/mL) reduction in bacterial population was observed for R2-K1 and R2-K11, respectively. Additionally, all four capsule-targeting R2 tailocins showed a significant impact on the growth curve at 8 h. These observations emphasize again that this assay most easily captures functional tailocins, even when having inferior activity necessitating further optimization.

**FIG 6.**
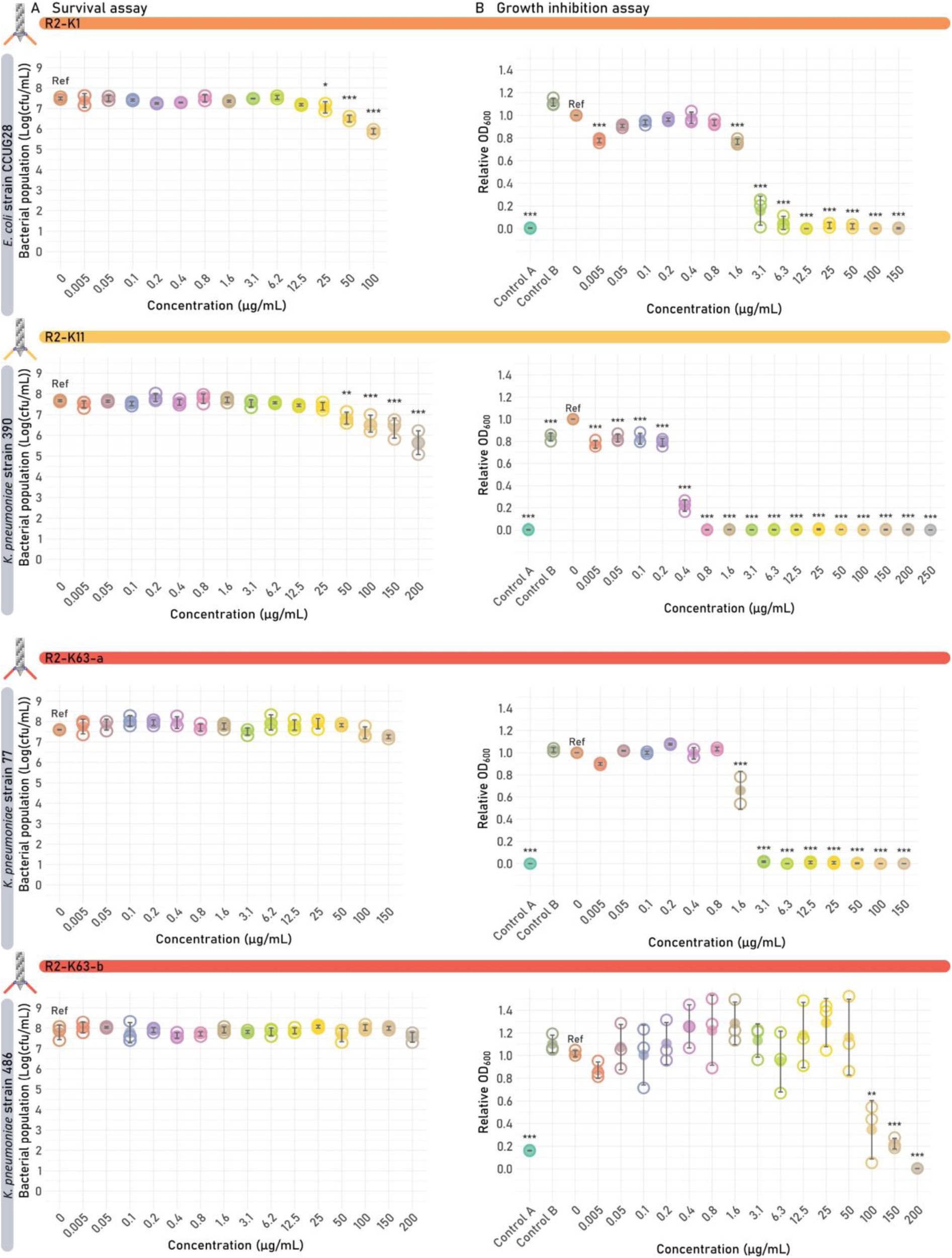
Survival and growth inhibition assays of the capsule-targeting R2 tailocins. Each R2 tailocin was tested on its susceptible *E. coli* and *K. pneumoniae* target strain. Significant differences are shown by asterisks (*p < 0.05; **p < 0.005, ***p < 0.001). (**A**) Survival assay. One plot is shown for each R2 tailocin construct, showing the bacterial colony count in function of the concentration of the added R2 tailocin. The value of each biological replicate is displayed using open circles and the mean values are shown as full circles. (**B**) Results of the growth inhibition assay at 8 h are shown per R2 tailocin construct. The relative OD_600_ of each biological replicate is displayed using open circles, and the mean relative OD_600_ is shown as full circles. Two additional controls were performed, one containing R2 tailocin but lacking the bacterial strain (Control A) and one containing an RBP lacking mutant R2 tailocin particle (R2Δprf15) instead of the engineered R2 tailocin of interest (Control B). Both controls were added at the same concentration as the highest available R2 tailocin concentration (100-200 µg/mL).

It must be noted that the R2 tailocin concentration at which a bactericidal effect is observed is dependent on the purification method. Tailocin R2-K11 was produced using two different methods. This tailocin showed a significant bactericidal effect in the survival assay from a concentration of 50 µg/mL after purification by high-speed centrifugation but from 6.2 µg/mL after purification by AS precipitation. Similarly, at a R2 tailocin concentration of 100 µg/mL in the survival assay, an increase in activity from 1.1 ± 0.4 up to 6.3 ± 1.3 log(CFU/mL) was observed for R2-K11 when AS precipitation was the chosen protein purification method. Nevertheless, the concentration of purified tailocins obtained by AS precipitation was 1.6-fold lower than high-speed centrifugation, measured using the micro-BCA protein assay (**Supplementary FIG S2** and **S3**). A similar but less pronounced trend was observed for the MIC value, which was 2-fold lower when R2-K11 was purified by AS precipitation (**Supplementary FIG S1**). These observations indicate that the purity and/or intactness of the tailocins is higher in the case of AS precipitation. However, high-speed centrifugation excels in convenience.

### Construction of a bivalent R2 tailocin R2-K11-K1 with activity against both *K. pneumoniae* capsular serotype K11 and *E. coli* capsular serotype K1

The VersaTile DNA assembly method is dedicated to the rapid combinatorial construction of modular proteins and allows to pursue the construction and evaluation of more advanced RBP architectures in engineered tailocins. As explained for Klebsiella phage K11, many phages have a dual RBP architecture with the first RBP attached via its anchor domain to the phage tail. A second RBP then docks via its N-terminal conserved peptide (CP) on a branching domain present in the first RBP, which is located directly after the anchor domain. Specifically in phage K11, the first RBP of phage K11 (K11gp17) with K11 specificity (RBD-K11) is connected to the second RBP of phage K11 (K11gp43) with K31 specificity via such branching domain-CP connection.

To transplant such a dual RBP architecture into the R2 tailocin scaffold, the RBP cluster was built by assembling four tiles sourced from different tailocins or phages: (I) the anchor tile sourced from the native R2 tailocin; (II) RBD-K11 of Klebsiella phage K11 targeting the Klebsiella capsular serotype K11. RBD-K11 includes the T4gp10-like branching domain that serves as a crucial docking site for the second RBP; (III) the CP tile from the second RBP, KP32gp38, of Klebsiella phage KP32 (27), which is highly similar to its equivalent in K11 (26), and (IV) RBD-K1 from *E. coli* phage K1F targeting the *E. coli* capsular serotype K1 (**FIG 7**, a and b). This VersaTile assembled fragment encoding the dual RBP cluster was subsequently expressed *in trans* as described above. Similar to R2-K11 and R2-K1, which were produced simultaneously with R2-K11-K1, the R2-K11-K1 tailocin was purified by high-speed centrifugation and spotted onto the panel of *E. coli* and *K. pneumoniae* strains (**FIG 5**). R2-K11 showed clearance on *K. pneumoniae* strain 390 (K11 capsule), while R2-K1 showed clearance on *E. coli* strains CUGG28 (capsular serotype K1), CAB1 (capsular serotype K1) and ECOR28 (serogroup O104, capsular serotype unknown), with no cross-activity observed. In contrast, R2-K11-K1 displayed a broadened host range targeting both capsular serotypes, visible as opaque spots. The bactericidal activity of the bivalent tailocin was further evaluated against its susceptible hosts *K. pneumoniae* strain 390 (capsular serotype K11) and *E. coli* strain CUGG28 (capsular serotype K1), using the survival assay. A significant bactericidal effect was observed against both strains starting from 50 µg/mL (**FIG 7**, c). The activity of R2-K11-K1 at a concentration of 100 µg/mL in the survival assay was similar to the activity of R2-K1 and R2-K11, with a 1.4-2.0 log(CFU/mL) reduction in bacterial population (**TABLE 1**). A significant impact on the growth curve after 8 h could be observed from 0.8 µg/ml and 25 µg/mL for *K. pneumoniae* capsular serotype K11 and *E. coli* capsular serotype K1, respectively (**FIG 7**, d). A significant MIC after 24 hours was only observed for *K. pneumoniae* strain 390 (50 µg/mL). The impact on the growth curve provoked by the second RBP targeting *E. coli* K1 capsule was thus less distinct than for *K. pneumoniae* K11 capsule by RBP1.

**FIG 7.**
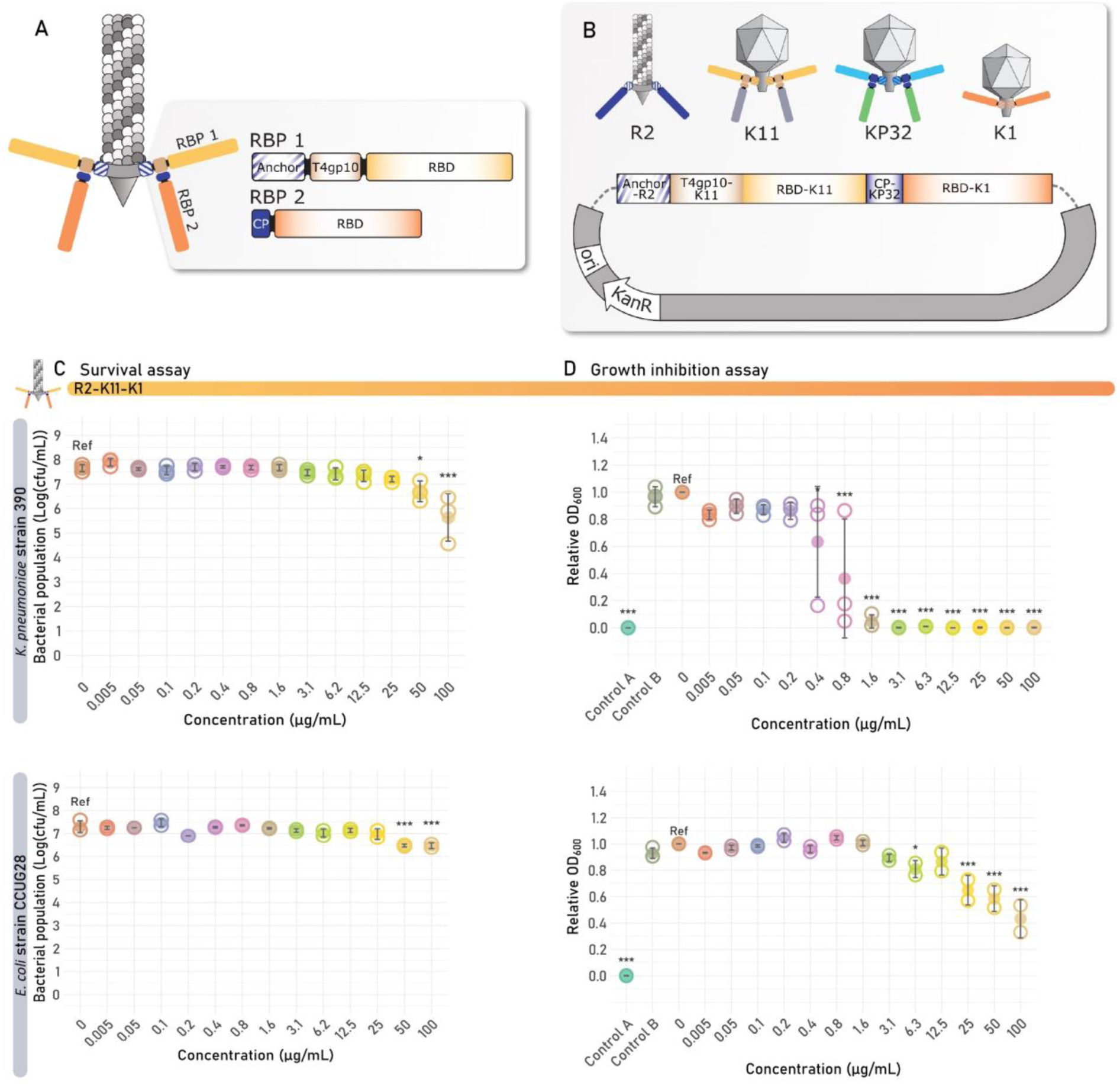
Analysis of bivalent R2 tailocin R2-K11-K1. (**A**) Modular build-up of R2 tailocin R2-K11-K1 with a dual receptor-binding protein (RBP) system with a branched anchor structure. The branched RBP system consists of four building blocks: (I) the N-terminal R2 anchor for attachment of the tailocin baseplate, (II) the RBD with specificity towards *Klebsiella* capsular serotype K11, sourced from phage K11gp17, including the T4gp10-like branching domain, (III) the conserved peptide (CP) sourced from phage KP3 to attach the second RBP to the branching domain of the first RBP, and (IV) the K1 RBD sourced from phage K1F. (**B**) The four tiles were assembled in the shuttle expression vector pVTD29 in a predefined order using VersaTile. (**C**) Survival assay. The bacterial colony count is shown in function of the concentration of the added R2 tailocin. The value of each biological replicate is displayed using open circles and the mean values are shown as filled circles. (d) Growth inhibition assay results at 8 h of the engineered bivalent R2 tailocin targeting both *K. pneumoniae* and *E. coli* capsular serotype. The relative OD_600_ of each biological replicate is displayed using open circles and the mean relative OD_600_ is shown as full circles. Two additional controls were performed, one containing R2 tailocin but lacking the bacterial strain (Control A) and one containing an RBP lacking mutant R2 tailocin particle (R2Δ*prf15*) (Control B). For both the survival and growth inhibition assays, the R2 tailocin was tested on its susceptible *E. coli* and *K. pneumoniae* target strains, which are indicated in vertical, gray-colored headings on the left side of the figure. Significant differences are shown by asterisks (*p < 0.05; **p < 0.005, ***p < 0.001).

## Discussion

In this work, we have converted the tailocin engineering and production method described previously (17) into a plug-and-play framework that facilitates the rapid and straightforward integration of RBPs from diverse origins. We have evaluated this tailocin platform with diverse RBPs targeting O-antigen or capsule (K-antigen) of *E. coli* and *K. pneumoniae*. In addition, we have demonstrated that also a branched RBP system targeting capsular serotypes of two different species can be transplanted into the R2 tailocin scaffold. In total, eight tailocins were redirected towards the O-antigen of *E. coli*, one towards the *E. coli* capsule, three towards the *K. pneumoniae* capsule, and one bivalent R2 tailocin targeting both the capsule of *K. pneumoniae* and *E. coli.* The main progress of this work compared to the state-of-the-art in tailocin engineering are the scale (i.e., the high number of engineered tailocins in a single study), the development of the first tailocins redirected towards the bacterial capsule and *K. pneumoniae,* and the first application of a branched RBP system in a tailocin scaffold.

### VersaTile applications to engineer tailocins

The enhancement in scale was achieved through the introduction of VersaTile (a DNA assembly technique dedicated to the combinatorial engineering of modular proteins such as RBPs) in the tailocin engineering process. Particularly the branched RBP system excels in modularity and is a prime example of how VersaTile can facilitate the rapid construction and exploration of modular RBP systems in a tailocin scaffold. This study expands on earlier work where the VersaTile technique was successfully implemented for other modular proteins such as phage lysins (23), innolysins (22), and cellulosomes (28). In addition, the VersaTile technique was successfully applied to create chimeric tailspikes by recombining the N-terminal anchor and C-terminal RBDs of four different Klebsiella phages (15). The efficiency of this assembly process can be attributed to the use of a single type IIs restriction enzyme that allows simultaneous restriction-ligation in contrast to traditional cumbersome cloning techniques that were used before for tailocin engineering. The VersaTile junction affected the antibacterial activity of R2 tailocins. However, traditional cloning methods also result in junctions without the flexibility to choose the amino acids and junction-less assembly lacks scalability because of the custom design process per engineered tailocin or synthesis cost (∼150€/construct). Upon identification of functionally engineered tailocins, these techniques can be used to further tune the hit-to-lead development and improve biological activity.

### R2 tailocin as the scaffold for enzymatic tailspikes and extra-species RBPs

An array of engineered R2 tailocins with a chimeric RBP were successfully created targeting the *E. coli* O-antigen serogroups O26, O103, O104, O111, O145, O146, and O157, *E. coli* capsular serotype K1 and *K. pneumoniae* capsular serotype K11 and K63. The majority of the newly engineered R2 tailocins, including R2-O104 and R2-O157, R2-O26, R2-O103-b, R2-O111, R2-O145-b, and R2-O146, R2-K1 and R2-K11 showed clearance in the spot assay, bactericidal activity in the survival assay, and a significant impact on the growth curve in the growth inhibition assay against their susceptible host(s). We proved that the *P. aeruginosa* R2 tailocin can be used as a scaffold to integrate extra-species RBPs with O-antigen and capsule specificity, showing activity against the respective serotype/host. Notably, the used RBPs in this work have a tailspike morphology featured by a β-helix in contrast to the fibrous tail fiber morphology of the native R2 RBP. Correct delineation of the RBD and optimization of the junction site are major aspects impacting the optimal functioning of newly engineered tailocins and should be the initial focus for further tailoring.

### R2 tailocin as the scaffold for complex RBP systems enabling cross-species activity

The first bivalent R2 tailocin R2-K11-K1 with activity against both the capsular serotype K11 of *K. pneumoniae* and capsular serotype K1 of *E. coli* emphasized the surprising engineering potential of the tailocin scaffold, here facilitated by the flexibility of VersaTile and the availability of a diverse repository of tiles of RBP modules. The chimeric RBP cluster with bivalent host specificity was created by reusing tiles from this in-house existing tile repository, which contains non-homologous, modular RBP building blocks sourced from different Escherichia and Klebsiella phages (**FIG 7**). Latka et al. already demonstrated that the anchor domain of Klebsiella phage KP32 could be fused with the enzymatic domain of RBP1 of Klebsiella phage K11 and successfully integrated this chimeric RBP in the phage K11 scaffold (15). This replacement was feasible as both phages belong to the same taxonomic genus and have a high identity of 88% at the amino acid level between their anchor domains (15, 26). Combining these findings with other unpublished data, an inter-species branched RBP cluster was created consisting of the RBD of Klebsiella phage K11gp17 with K11 capsule specificity in the first position and the RBD of Escherichia phage K1Fgp17 with K1 capsule specificity in the second position. The anchor-branched system relied on the T4gp10-like domain present in the Klebsiella phage K11gp17 and the CP derived from Klebsiella phage KP32gp38. Subsequently, the cluster was attached to the R2 tailocin scaffold using the native R2 anchor domain. In total, building blocks derived from four different origins (Pseudomonas, Escherichia, and two different Klebsiella phages) were used to create the first cross-species bivalent tailocin R2-K11-K1.

### Technical aspects of engineered tailocins production impacting the final antibacterial effectiveness

The number of R2 tailocin particles in the survival assay can be estimated with a formula based on the Poisson distribution (17). This number is relevant to calculate the tailocin-to-cell ratio, as tailocins are single-hit molecules and multiple tailocin particles may be randomly attached to a single bacterial cell, or conversely, some bacterial cells may not encounter any tailocin at all (29). However, the efficacy of the engineered R2 tailocins on each strain varies, meaning that this estimate on the number of killing particles would become strain-dependent. We therefore chose to assess the R2 tailocin efficacy based on protein concentration, as performed for traditional antimicrobial assays. This approach was also applied by Redero and colleagues when testing the activity of wild-type R-type tailocins in a murine pneumonia model (30). A downside of this method is that the R2 tailocin is not entirely pure after a single step purification, resulting in an underestimate of the tailocin particle number.

Tailocin purification by AS precipitation (17, 22, 29), PEG precipitation (30–32), ultra-centrifugal filter units (22), and/or high-speed centrifugation (17, 18, 20, 29, 30, 32) and combinations thereof have been reported. In the course of this study, we switched from AS precipitation to high-speed centrifugation due to its faster processing time and its higher amenability for upscaling, although the latter method resulted in a 16-fold lower bactericidal activity in the survival assay for R2-K11 (**Supplementary FIG S3**). Thus, R2 tailocin purification using high-speed centrifugation or AS purification alone can provide an initial indication of the tailocin activity, but a higher tailocin purity is needed to assess the activity in greater detail. Combining these purification methods with other purification methods such as chromatographic methods (33) or density gradient centrifugation (34) will increase the purity of the tailocins. In addition, protein concentration does not inform on the intactness and completeness of tailocin particles. An imbalance of the produced chimeric RBPs in relation to the available R2 scaffolds may lead to dysfunctional, partially assembled, or misassembled particles. In addition, it must be noted that F-type tailocins produced under the same *recA* regulator are possibly still present in the purified engineered R2 tailocins preparation. Removal of unnecessary genes under the *recA* regulator can be performed to improve the activity determination of tailocin product.

To easily detect newly engineered, functional tailocins, we found that assessment of growth curve after 8 h in the growth inhibition assay proved to be the most suitable method with a statistically significant impact for all engineered tailocins. Interestingly, statistically significant impact on the growth curve of R2-O157 was observed on target *E. coli* O157 strain 264, although there were no clear zones visible when spotting on bacterial lawns, underlining the relevance of the growth inhibition assay for initial testing, even if the spot assay is negative. However, whereas R2-O26, R2-K63-a, and R2-K63-b showed a significant impact on the growth curve, no bactericidal effect could be observed under the tested conditions. These observations confirm that the growth inhibition assay is more sensitive than other assays, which can be attributed to the higher tailocin-to-target cell ratio.

Interestingly, the switch from shuttle expression vector pVTD29 to pVTD27 for two tailocin constructs R2-O103-a and O145-a to obtain R2-O103-b and R2-O145-b, respectively, increased the antibacterial activity of the engineered tailocins. When shuttle expression vector pVTD27 without *lac* repressor was used to create R2-O103-b and R2-O145-b, a 16-and 540-fold improvement could be observed in the spot assay (**FIG 3**). Additionally, these -b variants performed 20 and 125 times better than the -a variants in the survival assay and 1260 and 620 times better in the growth inhibition assay at 8 h for R2-O103 and O145, respectively (**TABLE 1**; **FIG 4**). These results indicate that an increased *in trans* expression and possible accumulation of RBPs prior to induction of the R2 tailocin scaffold (due to the absence of the *lac* repressor) can significantly improve the quantity and proportion of completely assembled R2 tailocins. Thus, the amount of RBP may be the limiting factor for the assembly of intact tailocins. This increased RBP production may lead to more correctly or completely assembled tailocin particles, consequently resulting in lower required doses to exert a significant killing or growth inhibitory effect. To further investigate this hypothesis, the utilization of a shuttle expression vector featuring a tunable promotor capable of regulating gene expression levels (e.g. an arabinose-inducible promotor) could be explored. Alternatively, a PCR amplicon or synthetically ordered fragment of the engineered RBP can be provided *in vitro* into a cell-free system to produce tailocins (11).

### Tailocins, RBPs, and phages as therapeutic and diagnostic tools

Tailocins, phages and enzymatic RBPs all have antibacterial properties but differ in their mode of action. Tailocins are non-replicative proteins that rely on a single-hit mechanism, binding to and puncturing the cell envelope to kill the cell. In contrast, phages are replicative entities that, after receptor recognition, inject their genetic material into the host, initiating a replication cycle that ultimately leads to bacterial lysis and cell death. Phages have a so-called auto-dosing effect of continued replication as long as new hosts can be infected. Yet, replication is also associated with possible evolutionary changes in the replicated phage genome. Enzymatic RBPs, on the other hand, do not directly kill bacterial cells but actively degrade surface polysaccharides, making the bacterium again more susceptible to immune system clearance of other conventional therapeutics (35). Tailocins and RBPs have more resemblance to conventional pharmaceuticals, thereby circumventing regulatory, ethical and legislative difficulties related to the use of phages.

One important factor to consider is the difference in activity range between phages and tailocins. Two observations can be made on the difference between the efficacy of R2 tailocins and phages in this work. First, Escherichia phage Tp10 containing the RBD that was used to create the chimeric RBP in R2-O157, infects only one out of six *E.coli* strains of the respective serogroup, namely strain Sakai (36). However, R2-O157 kills five out of six *E.coli* strains (332, 584, 777/1, 2905 and Sakai) with an additional impact on the growth curve for *E. coli* strain 264 (**FIG 4**). This result of R2-O157 suggests a broader activity range of R2 tailocins compared to phages. This can be explained by the presence of intracellular anti-viral defenses restricting the host spectrum of phages but not of tailocins. Secondly, Escherichia phage PAS7, delivering the RBD for tailocin R2-O103, infects *E.coli* strain 4215/4 and causes lysis from without on *E.coli* strains NVH-848 and P11-2315. This lytic activity on all three *E. coli* O103 strains suggests that Escherichia phage PAS7 can successfully bind to its host (37). In contrast, the engineered tailocin R2-O103 kills *E.coli* strains 4215/4 and NVH-848, but not P11-2315. Additionally, some other strains in this work are resistant to R2 tailocins, although having the predicted serogroup/capsular serotype. This variable susceptibility of different strains towards a single tailocin may be explained by other confounding factors such as possible masking of the O-antigen.

Based on the high RBP specificity, the engineered R2 tailocins could potentially also be applied as diagnostics for bacterial typing instead of phages, avoiding the bias introduced by phage defense systems. Tailocins may be superior to RBPs for diagnostic purposes since the tailocin-induced lysis effect is easier to detect compared to the halo zone produced by enzymatic RBPs.

## Materials and methods

### Media, bacterial strains, plasmids, and bacteriophages

**Supplementary TABLE S1** lists all bacterial strains and phages used in this study, including their source. *K. pneumoniae* and *P. aeruginosa* strains were cultured in Tryptic Soy Broth (TSB; Oxoid, Basingstoke, UK) and on Tryptic Soy Agar (TSA; Oxoid) at 37 °C. *E. coli* strains were grown in Lysogeny Broth (LB; 1% (*w/v*) tryptone, VWR, Radnor, PA, USA; 0.5% (*w/v*) yeast extract, VWR; 1% (*w/v*) NaCl, Fisher Scientific, Thermo Fisher Scientific, Waltham, MA) and on LB agar (LA; LB supplemented with 1.5% (*w/v*) bacteriological agar, VWR) at 37 °C. Soft agar used in spotting assays contained 0.5% (*w/v*) agar (VWR). Each tile from the tile repository (**FIG 1**) was initially cloned and stored in vector pVTE (**Supplementary FIG S4**), as previously described (23). *E. coli* TOP10 cells (Invitrogen, Carlsbad, CA, USA) transformed with cloning vector pVTE were grown on LB supplemented with 100 μg/mL ampicillin (Fisher Scientific, Thermo Fisher Scientific, Waltham, MA) and 5% (*w*/*v*) sucrose (Fisher Scientific). For the assembly and storage of the engineered tailocin RBP construct, pVTD27 or pVTD29, two modified versions of the pUCP*tac* expression vector (17) were created (**Supplementary FIG S5 and S6**). A *sacB* coding sequence, flanked with two BsaI recognition sites and two distinct six-nucleotide-long position tags (start (P_start_) and end (P_end_)) into which the BsaI restriction sites were embedded, was inserted downstream of the *tac* promotor. As a result, the *sacB* gene along with the BsaI recognition sites are removed during the VersaTile assembly reaction and replaced by the assembled RBP construct. Gentamicin (Carl Roth, Karlsruhe, Germany) and sucrose (Fisher Scientific) at final concentrations of 20 μg/mL and 5% (*w*/*v*), respectively, were supplemented to the media for *E. coli* TOP10 (Invitrogen) or *P. aeruginosa* to be transformed with expression vector pVTD27 or pVTD29 (**Supplementary FIG S5 and S6**).

### Engineering of the RBP gene

To create chimeric RBPs, anchor and RBD nucleotide sequence tiles were constructed. The anchor tile, derived from the native R2 tailocin, was created to facilitate the integration of different C-terminal RBD tiles from *E. coli* and *K. pneumoniae* phages in the R2 tailocin scaffold. Fragments of interest were amplified with Phusion High-Fidelity DNA Polymerase (Thermo Fisher Scientific, Waltham, MA) using VersaTile-specific primers featuring extended 5’ sequences to create the different tiles (**Supplementary TABLE S2 and S3**). The subsequent amplicons were verified with gel electrophoresis on a 1.2% (*w/v*) agarose (Chem-Lab, Zedelgem, Belgium) gel and purified with the GeneJET Gel Extraction Kit (Thermo Fisher Scientific). These amplicons were inserted in the pVTE vector by restriction-ligation (SapI; Thermo Fisher Scientific;T4 DNA ligase; Life Technologies, Carlsbad, CA, USA) (**Supplementary TABLE S4**). Subsequently, the anchor and RBD tiles were fused in the shuttle expression vector pVTD29 in the VersaTile assembly reaction (**Supplementary TABLE S5**). Two tailocin variants were created in both pVTD27 and pVTD29, which are identical but the former lacks the *lac* repressor (*lacI* gene). Cloning in pVTD27 results in the presence of a nine instead of six-nucleotide junction between the anchor and the RBD (encoding Leu-Gly-Ser vs Gly-Ser) resulting from the original vector backbone (pM50 (22) and the assembly junction site. All transformations in the VersaTile cloning and assembly steps are performed using chemically competent *E. coli* TOP10 cells (Invitrogen). Plasmid extraction was performed using the GeneJet plasmid miniprep Kit (Thermo Fisher Scientific). Overview figures were created using Affinity Designer 2 version 2.4.2.

### R2 tailocin production and purification

R2 tailocin expression and purification were based on previous research performed (17). Briefly, 18 h cultures of *P. aeruginosa* PAO1 *Δprf15* strains containing the pVTD27 or pVTD29 expression vector (depending on the construct, **Supplementary TABLE S6**) containing the engineered RBP coding sequence *in trans,* were grown and 1:100 diluted in 15 mL G medium (20 g/L monosodium L-glutamic acid (Sigma, St. Louis, MO, USA), 5 g/L glucose (Carl Roth), 2.67 g/L Na_2_HPO_4_.2H_2_O (Fisher Bioreagents, Thermo Fisher Scientific, Waltham, MA, USA), 100 mg/L MgSO_4_.7H_2_O (VWR), 250 mg/L KH_2_PO_4_ (Carl Roth), 500 mg yeast extract (VWR); pH 7.2) supplemented with 20 µg/mL gentamicin (Carl Roth) and incubated at 37 °C while shaking at 225 rpm. At an OD_600_ of 0.25, 3 µg/mL mitomycin C (Sigma) was added to induce the tailocin genes in the *P. aeruginosa* PAO1 *Δprf15* genome. When a RBP was expressed *in trans*, the cultures were additionally supplemented with 0.25 mM isopropyl β-D-1-thiogalactopyranoside (IPTG; Carl Roth) to induce the *tac* promotor. Thereafter, the culture was incubated (37 °C, 180 rpm) for 2.5 h until complete lysis was observed. Subsequently, 0.1 U/mL DNaseI (Thermo Fisher Scientific) was added, and the culture was additionally incubated for 30 min. After incubation, the cultures were centrifuged (Eppendorf 5810R; Eppendorf, Hamburg, Germany) at 20,000 × *g* for 1 h at 4 °C to remove the bacterial debris. Two methods of tailocin purification were performed (**Supplementary TABLE S6**). (I) A 4 M ammonium sulfate (AS, Carl Roth) solution was dropped at a slow flow rate of 1 mL/min to the supernatant while shaking on ice until a final concentration of 1.6 M was reached. The resulting suspensions from each replicate were thereafter stored for 18 h at 4 °C while shaking at 160 rpm. The AS concentrate was precipitated by centrifugation (Eppendorf 5810R) at 20,000 × *g* for 1 h at 4 °C and resuspended in 10% (*v/v*) of the start volume with cold TN50 buffer (1.21 g/L Tris-HCl (VWR), 2.92 g/L NaCl (Thermo Fisher Scientific); pH 7.5). (II) Alternative to AS precipitation, tailocins were precipitated using a high-speed centrifuge (Sorvall Lynx 6000, Thermo Fisher Scientific) at 80,000 × *g* for 1 h at 4 °C and resuspended in 10% (*v/v*) of the start volume with cold TN50 buffer (18). Each tailocin solution was filter sterilized with a 0.45 µm PES syringe filter (Novolab, Geraardsbergen, Belgium) to remove residual impurities. To evaluate the production host, the following three controls were included: (1) native R2 tailocin produced in *P. aeruginosa* PAO1 strain (R2-WT), (2) *P. aeruginosa* PAO1 *Δprf15* strain containing the pVTD29 expression vector encoding the native *prf15/16* gene *in trans* (R2-WT-trans), and (3) *P. aeruginosa* PAO1 *Δprf15* strain containing the pVTD29 expression vector encoding the VersaTile assembled *prf15/16* coding sequence *in trans* (R2-WT-VT). All expressions were performed in triplicate. An estimation of the tailocin concentration was made using the Micro BCA^TM^ Protein Assay Kit (Thermo Fisher Scientific) following the manufacturer’s instructions.

### Determination of the specificity spectrum

After tailocin expression, the specificity spectrum of the engineered R2 tailocin was determined by two spot assays. First, for each of the three replicates of engineered R2 tailocin production, an 18 h culture was grown for the four *K. pneumoniae* strains or twenty *E. coli* strains, and *P. aeruginosa* wtb/CF510, in triplicate. Next, a bacterial lawn was prepared from each culture using the soft agar method (38). This was achieved by adding 200 µL of the culture to 4 mL of molten soft agar that was subsequently poured onto a TSA/LA plate and left to dry. A 5 µL drop of the highest tailocin concentration was spotted on all bacterial lawns. Plates were then incubated for 18 h at 37 °C. Bacterial susceptibility became visible by the appearance of a clear, circular zone on the spot site. For comparison purposes, phages present in our collection, specifically Escherichia phages PAS7 (GenBank accession: OQ921331.1), PAS61 (GenBank accession: OQ921333.1), O157 typing phage 10 (Tp10; GenBank accession: KP869108.1), K1F (GenBank accession: NC_007636.1), Klebsiella phages K11 (GenBank accession: NC_011043), KP34 (GenBank accession: NC_013649.2), and KP36 (GenBank accession: NC_029099.1), from which the RBDs were transferred to the R2 tailocin scaffold, were also spotted on the bacterial strains containing the targeted receptor.

Once the susceptible host(s) were identified, an additional two-fold serial dilution of tailocins in TN50 buffer was made starting from a concentration of 300 µg/mL. Next, 3 µL of each dilution was spotted on a bacterial lawn of the tailocin-sensitive host prepared according to the soft agar method and incubated for 18 h at 37 °C. The minimal concentration was determined, for which the tailocin gave an opaque spot. TN50 buffer, the native R2 tailocin, and the RBP-deficient derivative (R2*Δprf15)*, also termed the tailocin scaffold, were spotted as negative controls.

### Survival assay to quantify the bactericidal killing effect

The culture of the target bacterial strain was grown for 18 h, diluted (1:100) in TSB/LB, and incubated at 37 °C (225 rpm) until the mid-exponential phase (OD_600_ = 0.6-0.7) was reached. Next, the bacteria were harvested by centrifugation (4000 × *g*, 10 min) and resuspended in 1:1 volume TN50 buffer, followed by an additional washing step. The following dilution series was made in TN50 buffer for the expressed tailocin: 150, 100, 50, 25, 12.5, 6.25, 3.13, 1.56, 0.78, 0.39, 0.19, 0. 0.097, 0.046 and 0 µg/mL. To start the assay, a volume of 50 µL of each tailocin dilution was added to 50 µL target bacteria. After 40 min of incubation at 37 °C, a ten-fold serial dilution of bacteria-tailocin mixture was made and 5 µL of the dilution series was spotted on a TSA/LA plate. After 18 h incubation at 37 °C, the colonies were counted and expressed in log_10_ scale (CFU/mL) and compared to the untreated control as an indication of tailocin bactericidal activity. The survival assay was performed for each of the three biological replicates. To test which tailocin treatments were significant across various concentration levels, a linear mixed model with random intercept per biological replicate was fitted to the data. Tailocin concentrations were considered as a categorical predictor. For each concentration, the outcome was compared to the untreated sample (tailocin concentration zero), while automatically adjusting for multiple testing using Dunnett’s approach. The impact of the different tailocin concentrations on the bacterial log-reduction compared to the untreated sample, termed the contrast, was plotted in relation to the tailocin concentrations with joint confidence intervals (**Supplementary FIG S7**) (39). The p-values were calculated to indicate the statistically significant differences in the results. Statistical analysis and data processing were performed in R Statistical Software ((40), R 4.3.1). Figures were further processed using Adobe Illustrator version 25.4.1 and Adobe InDesign version 16.4.3.

### Growth inhibition assay to assess the inhibitory effect of an engineered R2 tailocin against its sensitive bacterial strain(s)

Varying concentrations (250, 200, 150, 100, 50, 25, 12.5, 6.25, 3.13, 1.56, 0.78, 0.39, 0.19, 0.097, 0.046, and 0.0046 µg/mL) of the R2 tailocin were tested. For each independently produced tailocin, a dilution (1:100) of an 18 h incubated culture was prepared and grown until reaching an OD_600_ of 0.08-0.1. Subsequently, a dilution (1:100) in 2× TSB/LB was prepared, corresponding to a final bacteria density of ∼10^6^ CFU/mL. For each tailocin concentration, 50 µL of the bacterial suspension was added to 50 µL of the purified R2 tailocin derivative at the concentrations listed above. The optical density of the suspension was monitored at 600 nm using an Infinite 200 PRO® reader (Tecan, Männedorf, Switzerland) at 15-minute intervals over a 24-hour duration at 37 °C. The complete 24-hour monitoring period was visualized using R Statistical Software (v4.3.1, (Team, 2021)) and further processed using Adobe Illustrator version 25.4.1 and Adobe InDesign version 16.4.3. Negative controls consisted of (I) 50 µL of 2× TSB/LB added to 50 µL TN50 buffer (Blank), (II) 50 µL of 2× TSB/LB added to 50 µl R2 tailocin (highest available concentration) (control A), (III) 50 µL of the tailocin scaffold (R2*Δprf15*) in the same concentration as control A was added to 50 µL of bacterial suspension (control B), and (IV) 50 µL TN50 buffer added to 50 µL of bacterial suspension (concentration = 0 µg/mL).

The primary objective was to determine the concentration at which the tailocin completely inhibits bacterial growth. To achieve this, two different metrics were assessed based on the optical density measurements. As a first metric, a time point of approximately 8 h after exposure was chosen for all experiments based on the most substantial difference in inhibition of bacterial growth between the concentrations of the engineered R2 tailocin and the untreated sample. At this selected time point, the means of the relative optical densities were calculated for each concentration of tailocin tested and plotted against their corresponding tailocin concentrations. The relative OD_600_ represents a measure of bacterial growth, obtained by subtracting the baseline value (Blank) from all OD_600_ readings and then normalizing each value against the OD_600_ value of the untreated sample (tailocin concentration zero), ensuring that all values are within the range of 0 to 1. Statistics were performed using a Kruskal-Wallis test after failing assumptions, followed by a one-sided Dunnett’s test with the untreated sample as a reference. The resulting p-values were used to assess the significant differences with the untreated sample and were displayed in the figure presenting the results. All graphics were created using R Statistical Software ((40), R 4.3.1) and further processed using Adobe Illustrator version 25.4.1 and Adobe InDesign version 16.4.3. The second metric is the minimum inhibitory concentration (MIC), which corresponds to the lowest concentration at which full inhibition of growth was observed after 24 h. More specifically, full inhibition was assumed for the lowest concentration of R2 tailocin that inhibits ≥ 90% of the bacterial growth of the untreated sample at timepoint 24 h (41).

## Acknowledgments

We would like to thank Rani Dams for her graphical support. We are grateful to thank Dr. Dean Scholl of the AvidBiotics Corporation, and prof. Lone Brønsted of the Department of Veterinary and Animal Sciences of the University of Copenhagen (Denmark) for providing the R2 tailocin engineering plasmids. Additionally, we would like to express our gratitude to the Fostering Innovative Research based on Evidence (FIRE) at Ghent University (UGent) for their statistical consultancy services.

We would also like to thank Prof. Toril Lindbäck of the Department of Paraclinical Sciences at the University of Life Sciences in Oslo, Norway for providing us with *E. coli* strain NVH-848; Dr. Roger Stephan of the Institute for Food Safety and Hygiene of the University of Zurich, Switzerland, who provided most of the bacterial *E. coli* host strains; Prof. David Gally of the University of Edinburgh (Scotland) for kindly providing Escherichia O157 typing phage 10 (Tp10) and *E. coli* strain Sakai (Stx-); Prof. Ian J. Molineux of the Department of Molecular Biosciences, University of Texas at Austin, for kindly providing Escherichia phage K1F and *E. coli* strain CAB1; The Synthetic Biology Group of the Massachusetts Institute of Technology (USA) for kindly providing Klebsiella phage K11 and lastly, prof. Lars Jelsbak of the Department of Biotechnology and Biomedicine, Technical University of Denmark (Denmark) for kindly providing *P. aeruginosa* strain CF510.

C.P. and A.L. are supported by the Research Foundation—Flanders (FWO; 1S79422N; 1240021N; 1251224N). ZDK is supported by the National Science Center UMO-2022/47/I/NZ1/01450 and UMO-2022/04/Y/NZ6/00123 under the framework of the JPIAMR – Joint Programming Initiative on Antimicrobial Resistance.

## Ethics approval and consent to participate

Not applicable.

## Consent to publication

Not applicable.

## References

1. Koulenti D, Song A, Ellingboe A, Abdul-Aziz MH, Harris P, Gavey E, Lipman J. 2019. Infections by multidrug-resistant Gram-negative Bacteria: What’s new in our arsenal and what’s in the pipeline? Int J of Antimicrob Agents 53:211–224.

2. Naghavi M, Vollset SE, Ikuta KS, Swetschinski LR, Gray AP, Wool EE, Robles Aguilar G, Mestrovic T, Smith G, Han C, Hsu RL, Chalek J, Araki DT, Chung E, Raggi C, Gershberg Hayoon A, Davis Weaver N, Lindstedt PA, Smith AE, Altay U, Bhattacharjee NV, Giannakis K, Fell F, McManigal B, Ekapirat N, Mendes JA, Runghien T, Srimokla O, Abdelkader A, Abd-Elsalam S, Aboagye RG, Abolhassani H, Abualruz H, Abubakar U, Abukhadijah HJ, Aburuz S, Abu-Zaid A, Achalapong S, Addo IY, Adekanmbi V, Adeyeoluwa TE, Adnani QES, Adzigbli LA, Afzal MS, Afzal S, Agodi A, Ahlstrom AJ, Ahmad A, Ahmad S, Ahmad T, et al. 2024. Global burden of bacterial antimicrobial resistance 1990–2021: a systematic analysis with forecasts to 2050. Lancet doi:10.1016/s0140-6736(24)01867-1.

3. Ghequire MGK, De Mot R. 2015. The Tailocin Tale: Peeling off Phage Tails. Trends Microbiol 23:587–590.

4. Scholl D. 2017. Phage Tail-Like Bacteriocins, p 453–467. In Enquist L (ed), Annual Review of Virology, Vol 4, vol 4. Annual Reviews, Palo Alto.

5. Suleman M, Yaseen AR, Ahmed S, Khan Z, Irshad A, Pervaiz A, Rahman HH, Azhar M. 2024. Pyocins and Beyond: Exploring the World of Bacteriocins in *Pseudomonas aeruginosa*. Probiotics and Antimicrob Proteins doi:10.1007/s12602-024-10322-3.

6. Buth SA, Shneider MM, Scholl D, Leiman PG. 2018. Structure and Analysis of R1 and R2 Pyocin Receptor-Binding Fibers. Viruses 10:427.

7. Saha S, Ojobor CD, Li ASC, Mackinnon E, North OI, Bondy-Denomy J, Lam JS, Ensminger AW, Maxwell KL, Davidson AR. 2023. F-Type Pyocins Are Diverse Noncontractile Phage Tail-Like Weapons for Killing *Pseudomonas aeruginosa*. J Bacteriol 205:17.

8. Nobrega FL, Vlot M, de Jonge PA, Dreesens LL, Beaumont HJE, Lavigne R, Dutilh BE, Brouns SJJ. 2018. Targeting mechanisms of tailed bacteriophages. Nat Rev Microbiology 16:760–773.

9. Pirnay JP, De Vos D, Verbeken G, Merabishvili M, Chanishvili N, Vaneechoutte M, Zizi M, Laire G, Lavigne R, Huys I, Van den Mooter G, Buckling A, Debarbieux L, Pouillot F, Azeredo J, Kutter E, Dublanchet A, Górski A, Adamia R. 2011. The Phage Therapy Paradigm: *Prêt-à - Porter* or *Sur-mesure*? Pharm Res 28:934–937.

10. Pottie I, Vázquez Fernández R, Van De Wiele T, Briers Y. 2024. Phage lysins for intestinal microbiome modulation: current challenges and enabling techniques. Gut Microbes 16.

11. Woudstra C, Sorensen AN, Sorensen MCH, Brondsted L. 2024. Strategies for developing phages into novel antimicrobial tailocins. Trends Microbiol 10.1016/j.tim.2024.03.003:11.

12. Woudstra C, Sorensen AN, Brondsted L. 2023. Engineering of *Salmonella* Phages into Novel Antimicrobial Tailocins. Cells 12:15.

13. Yao GC, Le T, Korn AM, Peterson HN, Liu M, Gonzalez CF, Gill JJ. 2023. Phage Milagro: a platform for engineering a broad host range virulent phage for Burkholderia. J Virol doi:10.1128/jvi.00850-23:18.

14. Dams D, Brondsted L, Drulis-Kawa Z, Briers Y. 2019. Engineering of receptor-binding proteins in bacteriophages and phage tail-like bacteriocins. Biochem Soc Trans 47:449–460.

15. Latka A, Lemire S, Grimon D, Dams D, Maciejewska B, Lu T, Drulis-Kawa Z, Briers Y. 2021. Engineering the Modular Receptor-Binding Proteins of *Klebsiella* Phages Switches Their Capsule Serotype Specificity. mBio 12:12.

16. Lenneman BR, Fernbach J, Loessner MJ, Lu TK, Kilcher S. 2021. Enhancing phage therapy through synthetic biology and genome engineering. Curr Opin Biotechnol 68:151–159.

17. Williams SR, Gebhart D, Martin DW, Scholl D. 2008. Retargeting R-type pyocins to generate novel bactericidal protein complexes. Appl Environl Microbiol 74:3868–3876.

18. Ritchie JM, Greenwich JL, Davis BM, Bronson RT, Gebhart D, Williams SR, Martin D, Scholl D, Waldor MK. 2011. An *Escherichia coli* O157-Specific Engineered Pyocin Prevents and Ameliorates Infection by *E. coli* O157:H7 in an Animal Model of Diarrheal Disease. Antimicrob Agents Chemother 55:5469–5474.

19. Scholl D, Cooley M, Williams SR, Gebhart D, Martin D, Bates A, Mandrell R. 2009. An Engineered R-Type Pyocin Is a Highly Specific and Sensitive Bactericidal Agent for the Food-Borne Pathogen *Escherichia coli* O157:H7. Antimicrob Agents Chemother 53:3074–3080.

20. Scholl D, Gebhart D, Williams SR, Bates A, Mandrell R. 2012. Genome Sequence of *E. coli* O104:H4 Leads to Rapid Development of a Targeted Antimicrobial Agent against This Emerging Pathogen. PloS One 7:5.

21. Gebhart D, Williams SR, Scholl D. 2017. Bacteriophage SP6 encodes a second tailspike protein that recognizes Salmonella enterica serogroups O157:H7. Virology 507:263–266.

22. Zampara A, Sorensen MCH, Gencay YE, Grimon D, Kristiansen SH, Jorgensen LS, Kristensen JR, Briers Y, Elsser-Gravesen A, Brondsted L. 2021. Developing Innolysins Against *Campylobacter jejuni* Using a Novel Prophage Receptor-Binding Protein. Front Microbiol 12:13.

23. Gerstmans H, Grimon D, Gutiérrez D, Lood C, Rodríguez A, van Noort V, Lammertyn J, Lavigne R, Briers Y. 2020. A VersaTile-driven platform for rapid hit-to-lead development of engineered lysins. Sci Adv 6:11.

24. Pas C, Fieseler L, Pothier JF, Briers Y. 2024. Isolation, characterization, and receptor-binding protein specificity of phages PAS7, PAS59 and PAS61 infecting Shiga toxin-producing Escherichia coli O103 and O146 doi:10.21203/rs.3.rs-4758770/v1. Springer Science and Business Media LLC.

25. Arredondo-Alonso S, Blundell-Hunter G, Fu ZY, Gladstone RA, Fillol-Salom A, Loraine J, Cloutman-Green E, Johnsen PJ, Samuelsen O, Pöntinen AK, Cléon F, Chavez-Bueno S, De la Cruz MA, Ares MA, Vongsouvath M, Chmielarczyk A, Horner C, Klein N, McNally A, Reis JN, Penadés JR, Thomson NR, Corander J, Taylor PW, McCarthy AJ. 2023. Evolutionary and functional history of the *Escherichia coli* K1 capsule. Nat Commun 14:17.

26. Latka A, Leiman PG, Drulis-Kawa Z, Briers Y. 2019. Modeling the Architecture of Depolymerase-Containing Receptor Binding Proteins in *Klebsiella* Phages. Front Microbiol 10:20.

27. Squeglia F, Maciejewska B, Latka A, Ruggiero A, Briers Y, Drulis-Kawa Z, Berisio R. 2020. Structural and Functional Studies of a *Klebsiella* Phage Capsule Depolymerase Tailspike: Mechanistic Insights into Capsular Degradation. Structure 28:613-+.

28. Vanderstraeten J, Da Fonseca MJM, De Groote P, Grimon D, Gerstmans H, Kahn A, Moraïs S, Bayer EA, Briers Y. 2022. Combinatorial assembly and optimisation of designer cellulosomes: a galactomannan case study. Biotechnol Biofuels Bioprod 15.

29. Scholl D, Martin DW. 2008. Antibacterial efficacy of R-type pyocins towards *Pseudomonas aeruginosa* in a murine peritonitis model. Antimicrob Agents Chemother 52:1647–1652.

30. Redero M, Aznar J, Prieto AI. 2020. Antibacterial efficacy of R-type pyocins against *Pseudomonas aeruginosa* on biofilms and in a murine model of acute lung infection. Antimicrob Agents Chemother doi:10.1093/jac/dkaa121.

31. Baltrus DA, Clark M, Smith C, Hockett KL. 2019. Localized recombination drives diversification of killing spectra for phage-derived syringacins. Isme J 13:237–249.

32. Gebhart D, Lok S, Clare S, Tomas M, Stares M, Scholl D, Donskey CJ, Lawley TD, Govoni GR. 2015. A modified R-type bacteriocin specifically targeting Clostridium difficile prevents colonization of mice without affecting gut microbiota diversity. mBio 6.

33. Regulski K, Champion-Arnaud P, Gabard J. 2021. Bacteriophage Manufacturing: From Early Twentieth-Century Processes to Current GMP, p 699–729 doi:10.1007/978-3-319-41986-2_25. Springer International Publishing.

34. Lee FKN, Dudas KC, Hanson JA, Nelson MB, Loverde PT, Apicella MA. 1999. The R-Type Pyocin of *Pseudomonas aeruginosa* C Is a Bacteriophage Tail-Like Particle That Contains Single-Stranded DNA. Infect and Immun 67:717–725.

35. Abdelkader K, Gerstmans H, Saafan A, Dishisha T, Briers Y. 2019. The Preclinical and Clinical Progress of Bacteriophages and Their Lytic Enzymes: The Parts are Easier than the Whole. Viruses 11:96.

36. Dahan S, Knutton S, Shaw RK, Crepin VF, Dougan G, Frankel G. 2004. Transcriptome of Enterohemorrhagic *Escherichia coli* O157 Adhering to Eukaryotic Plasma Membranes. Infect and Immun 72:5452–5459.

37. Abedon ST. 2011. Lysis from without. Bacteriophage 1:46–49.

38. Kutter E. 2009. Phage host range and efficiency of plating. Methods Mol Biol 501:141–9.

39. Bates D, Mächler M, Bolker BM, Walker SC. 2015. Fitting Linear Mixed-Effects Models Using lme4. J Stat Softw 67:1–48.

40. Team RC. 2021. R: A language and environment for statistical computing. https://www.R-project.org/. Accessed

41. Tiwari S, Nizet O, Dillon N. 2023. Development of a high-throughput minimum inhibitory concentration (HT-MIC) testing workflow. Front Microbiol 14.

